# Landscape of human spinal cord cell type diversity at midgestation

**DOI:** 10.1101/2021.12.29.473693

**Authors:** Jimena Andersen, Nicholas Thom, Jennifer L. Shadrach, Xiaoyu Chen, Neal D. Amin, Se-Jin Yoon, William J. Greenleaf, Fabian Müller, Anca M. Pașca, Julia A. Kaltschmidt, Sergiu Pașca

**Affiliations:** Department of Psychiatry and Behavioral Sciences, Stanford University, USA; Stanford Brain Organogenesis, Wu Tsai Neurosciences Institute, USA; Department of Neurosurgery, Stanford University, USA; Department of Genetics, Stanford University, USA; Department of Applied Physics, Stanford University, USA; Chan–Zuckerberg Biohub, San Francisco, USA; Center for Bioinformatics, Saarland University, Germany; Department of Pediatrics, Division of Neonatology, Stanford University, USA

## Abstract

Understanding spinal cord generation and assembly is essential to elucidate how motor behavior is controlled and how disorders arise. The cellular landscape of the human spinal cord remains, however, insufficiently explored. Here, we profiled the midgestation human spinal cord with single cell-resolution and discovered, even at this fetal stage, remarkable heterogeneity across and within cell types. Glia displayed diversity related to positional identity along the dorso-ventral and rostro-caudal axes, while astrocytes with specialized transcriptional programs mapped onto distinct histological domains. We discovered a surprisingly early diversification of alpha (α) and gamma (γ) motor neurons that control and modulate contraction of muscle fibers, which was suggestive of accelerated developmental timing in human spinal cord compared to rodents. Together with mapping of disease-related genes, this transcriptional profile of the developing human spinal cord opens new avenues for interrogating the cellular basis of motor control and related disorders in humans.

## Introduction

The spinal cord plays a central role in integrating sensory and motor information to regulate movement. This is achieved by the coordinated function of motor neurons and other neuronal and glial populations that are arranged in spatially distinct domains (Côté et al., 2018; Haim and Rowitch, 2017; Jessell, 2000). Damage or degeneration of the spinal cord can lead to devastating disorders, such as amyotrophic lateral sclerosis (ALS) or developmental disorders, such as spinal muscular atrophy (SMA) or childhood leukodystrophies (Kanning et al., 2010; Monani, 2005; Swash et al., 1986). While cell diversity of the developing human neural tube has started to be unveiled (Rayon et al., 2021), no comprehensive profiling of the human spinal cord at midgestation is available. Access to this fetal stage of development is important as increasing evidence points at human-specific timing and characteristics in nervous system assembly (Gu et al., 2017; Oberheim et al., 2009; Silbereis et al., 2016; Sousa et al., 2017). For example, development of the cerebral cortex in humans takes place at a slower rate compared to rodents and other primates (Somel et al., 2009; Sousa et al., 2017). On the other hand, the human spinal cord has been proposed to develop and mature faster compared to rodents (Ohmura and Kuniyoshi, 2017; Tadros et al., 2015). Here, we used single cell transcriptomics to generate a cell type census and explore developmental landmarks in the fetal human spinal cord. We uncovered remarkable cellular diversity along several axes and discovered a surprisingly early diversification of alpha (α) and gamma (γ) motor neuron identity, highlighting differences in species-specific developmental timing.

## Results

### A transcriptional landscape of the developing human spinal cord

We used 10x Chromium to profile single cells and single nuclei from four samples at post-conception weeks (PCW) 17 and PCW18 (**Fig. 1A**, **S1A, B**; **Table S1**). To increase the probability of capturing motor neurons, we also performed Thy1-immunopanning or NeuN-sorting (**Fig. S1C**). Following quality control, doublet removal and filtering (performed separately in cells and nuclei), we obtained transcriptomes for 112,554 cells and 34,884 nuclei (**Fig. S1B, G**; **Table S2, S3**). PCW17 and PCW18 samples were highly correlated and were analyzed together (**Fig. S1H–J**). Single cell and single nucleus samples displayed some differences, as reported (Bakken et al., 2018). Specifically, we detected a higher number of molecules (nCount) and genes (nFeature) as well as a higher percentage of ribosomal and mitochondrial genes in cells, but a higher ratio of unspliced to spliced counts in nuclei (**Fig. S1G**). Differential expression and gene ontology (GO) analysis showed that nuclei-enriched genes were related to synapse organization (*NRXN1, SHANK1*) and cell adhesion (*CDH18, PCDH9*), while genes enriched in single cells were related to housekeeping (*GAPDH*), cellular stress (*DNAJA1, HSPB1*) and immediate early gene response (*FOS, JUN*), differences possibly associated with dissociation artifacts (**Fig. S2A–C**; **Table S4**).

**Figure 1:**
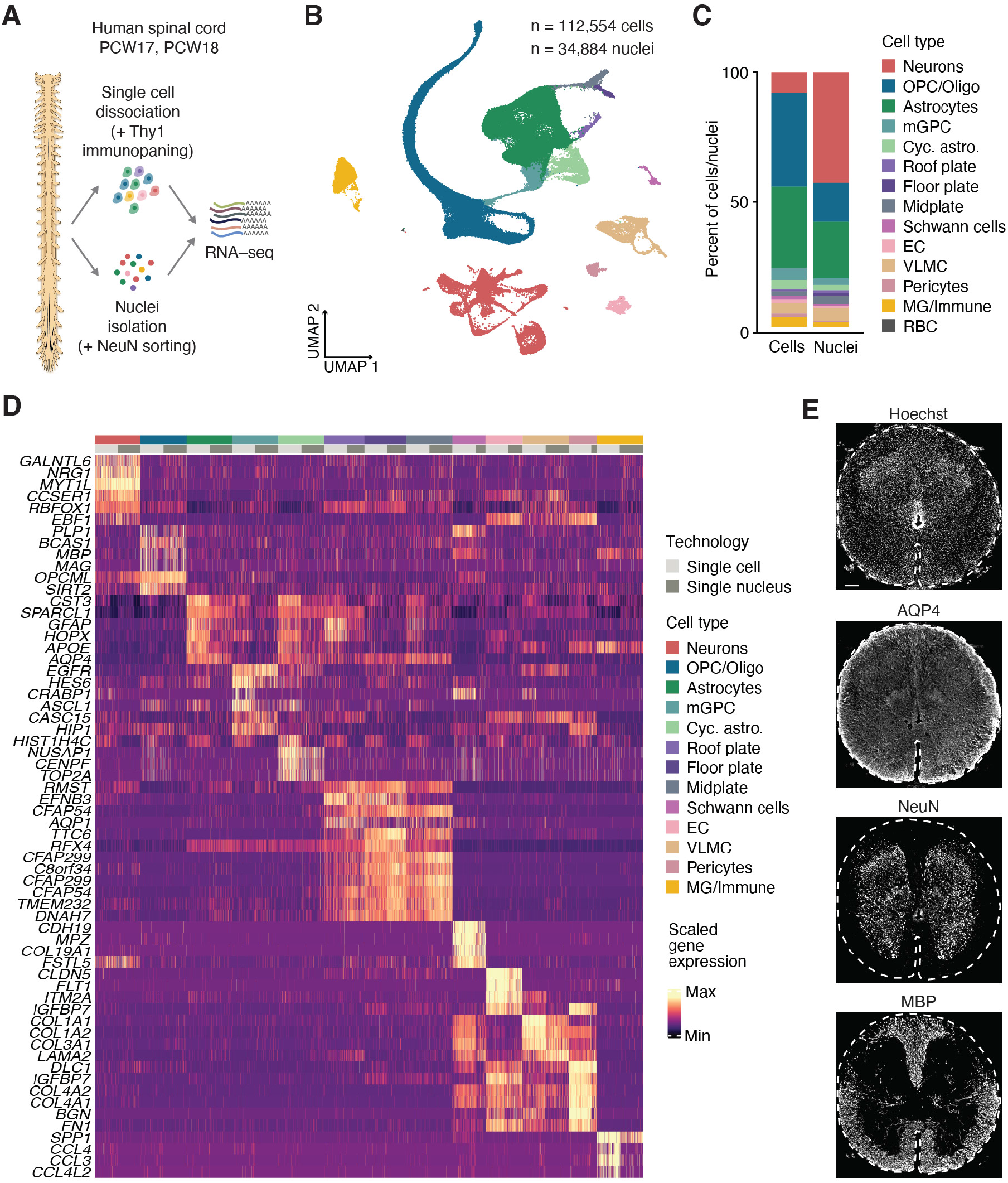
A single cell atlas of the human spinal cord at midgestation. **(A)** Schematic showing an overview of the samples and methods used for the study. PCW, postconception weeks. **(B)** UMAP representation of integrated single cells and single nuclei, colored by cell type. OPC/Oligo, oligodendrocyte precursor cell/oligodendrocyte; mGPC, multipotent glial progenitor cell; Cyc. astro., cycling astrocyte; EC, endothelial cell; VLMC, vascular leptomeningeal cell; MG/Immune, microglia/Immune; RBC, red blood cell. **(C)** Bar plot showing the percent of cell types in single cell and single nucleus samples. **(D)** Heatmap showing normalized expression of the top 4-6 unique markers per cell type shown separated by isolation method, and downsampled to at most 500 cells per cell type and isolation method. **(E)** Representative immunohistochemistry images of AQP4, NeuN (RBFOX3), and MBP in a coronal thoracic spinal cord cryosection at PCW19. Hoechst shows nuclei. Scale bar: 200 μm (**E**).

We used UMAP dimensionality reduction to visualize and cluster cells following the integration of single cells and single nuclei (Stuart et al., 2019)(**Fig. 1B**; **Table S5**). We found a group of astrocytes and cycling astrocytes as well as cells within the astroglia lineage (floor plate, roof plate, and midplate), cells in the oligodendrocyte lineage (OPC/Oligo), and a group of multipotent glial progenitor cells (mGPC) linking the two lineages. We also identified a group of neurons that was enriched in the single nucleus samples (**Fig. 1C**), and groups of vascular cells including endothelial cells (EC), pericytes and vascular leptomeningeal cells (VLMC), Schwann cells, and immune cells including microglia and monocytes (MG/Immune) (**Fig. 1D, E**). Cycling cells were distributed within the UMAP landscape and could be found as part of the OPC/Oligo, VLMC, EC, MG/Immune and Schwann cell groups (**Fig. S1K**).

### Pseudotime analysis revealed cellular processes enriched along the OPC/Oligo lineage

We first examined OPC/Oligo cells (**Fig. 2A**). Subclustering identified cell types along the oligodendrocyte lineage (**Fig. 2B**, **S3A–C**; **Table S6**): oligodendrocyte progenitor cells (OPCs), cycling OPCs and OPC-like mGPCs, differentiation-committed oligodendrocyte precursors (COP), newly formed oligodendrocytes (NFOL), myelin-forming oligodendrocytes (MFOL) and mature oligodendrocytes (MOL). Upon further inspection, we noticed that the OPC cluster included cells expressing genes associated with a dorsal identity (*PAX3, ZIC1*) and other cells expressing genes associated with ventral identity (*FLRT2, GABRB2*), presumably indicating that these progenitors were generated in specific spinal cord domains (**Fig. S3D**). Interestingly, this division was not apparent in the rest of the lineage.

**Figure 2:**
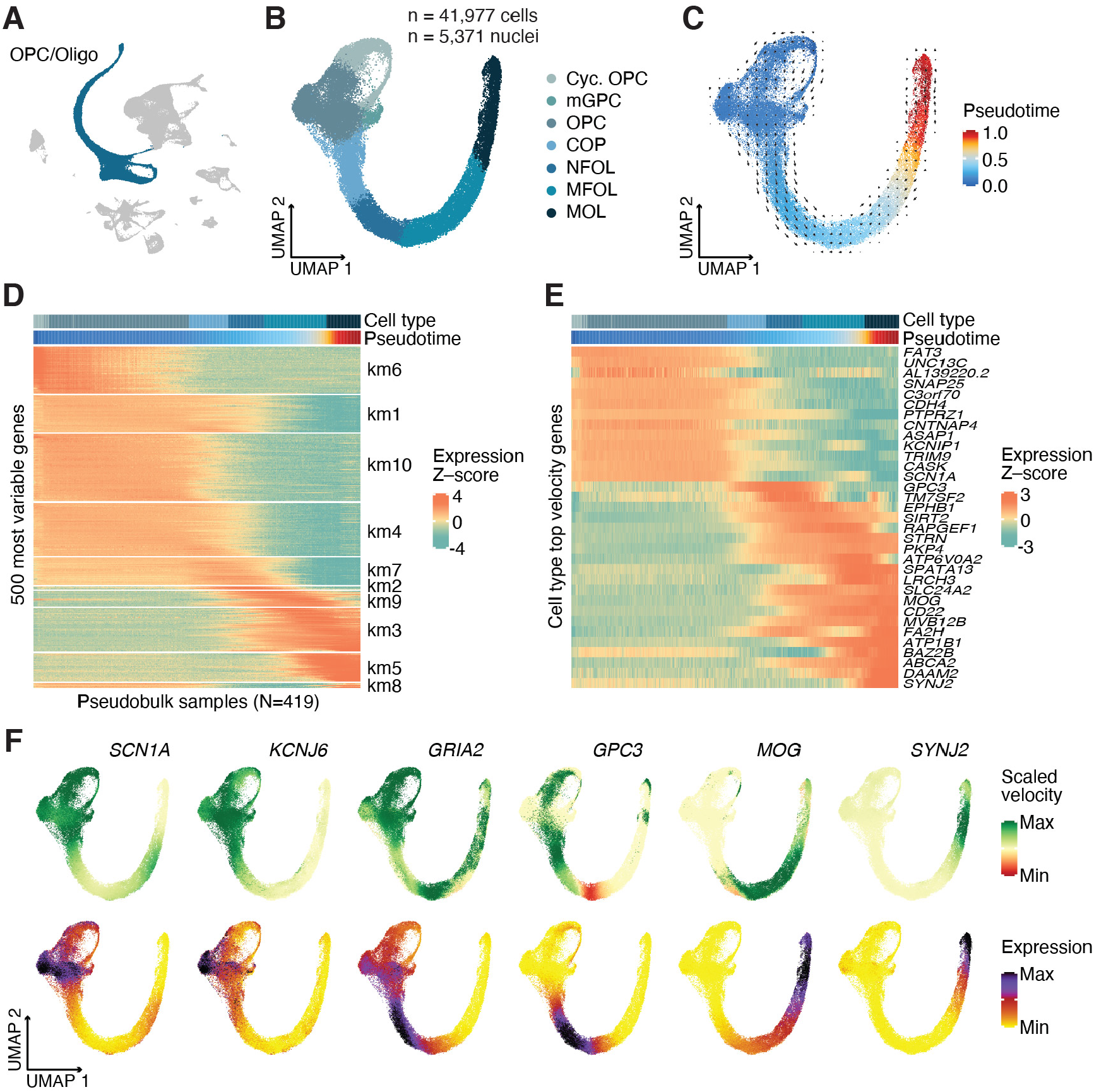
Oligodendrocyte lineage in the spinal cord. **(A)** Highlight of oligodendrocyte lineage cells in the main UMAP, showing the cells selected for subclustering. **(B)** UMAP of the OPC/Oligo subcluster colored by cell type. Cyc. OPC, cycling oligodendrocyte progenitor cells; mGPC, multipotent glial progenitor cells; COP, differentiation-committed oligodendrocyte precursors; NFOL, newly formed oligodendrocytes; MFOL, myelin-forming oligodendrocytes; MOL, mature oligodendrocytes. **(C)** UMAP showing the OPC/Oligo subcluster colored by the diffusion pseudotime using single-cell velocity as computed by scVelo. The arrow overlay shows aggregate projected velocities. **(D)** Heatmap showing scaled expression of the 500 most variable genes across 419 pseudobulk samples aggregated along pseudotime bins. Genes were clustered into 10 groups using k-means. **(E)** Heatmap showing scaled expression of cell-type dynamic genes. For each cell type the 5 highest-ranking genes according to velocity are shown. **(F)** UMAPs showing examples of dynamic genes with cell-type specific velocity along the trajectory showing computed velocity (top) and normalized expression (bottom).

To explore developmental progression in the OPC/Oligo lineage, we computed RNA velocity (Bergen et al., 2020; La Manno et al., 2018). Diffusion pseudotime analysis revealed putative origins of the trajectory in cycling cells (**Fig. S3E**) and inferred a differentiation trajectory. We annotated each cell with a pseudotime value (**Fig. 2C**) and grouped variable genes along this pseudotime into 10 clusters (**Fig. 2D**). GO analysis highlighted a sequence of cellular processes along the trajectory. Early in the pseudotime we found terms that included cell division, and terms associated with potassium transport and synapse assembly. Genes expressed in NFOL and MFOL were enriched for terms associated with axon guidance and migration, while MFOL and MOL were enriched for myelination-related genes (**Fig. S3E**). For each major cell type, we identified marker genes exhibiting particularly high estimated velocities, some of which manifested in increased expression downstream in the trajectory (**Fig. 2E, F**; **Table S7**). For instance, *GPC3* was characterized by high velocity in COPs and high expression in NFOLs followed by negative velocities indicating downregulation in MFOLs.

### Astrocytes in the human spinal cord show diversity along multiple axes

In the rodent spinal cord, astrocytes are spatially and functionally heterogeneous (Bayraktar et al., 2015; Haim and Rowitch, 2017; Hochstim et al., 2008; Khakh and Deneen, 2019; Molofsky et al., 2014; Tsai et al., 2012). To investigate this diversity in humans, we next subclustered astroglia (**Fig. 3A–C, S4A–C**; **Table S8**). We found a group of glial progenitors corresponding to ventricular zone cells and dorsal and ventral mGPCs that expressed *SOX9*, but lower levels of the more mature astrocyte markers *AQP4* and *SPARCL1* (**Fig. S4D**), cycling cells (**Fig. S4E, F**), and a group of cells we called “active” based on their expression of activitydependent genes such as *FOS*, *EGR1* and *ARC*. These cells were found almost exclusively in single cell samples (**Fig. S4A**) and their signature did not overlap with that of reactive astrocytes (**Fig. S4G**) (Liddelow et al., 2017), suggesting a signature likely related to the dissociation method. In addition to these, we found nine clusters of astrocytes along two axes of diversity.

**Figure 3:**
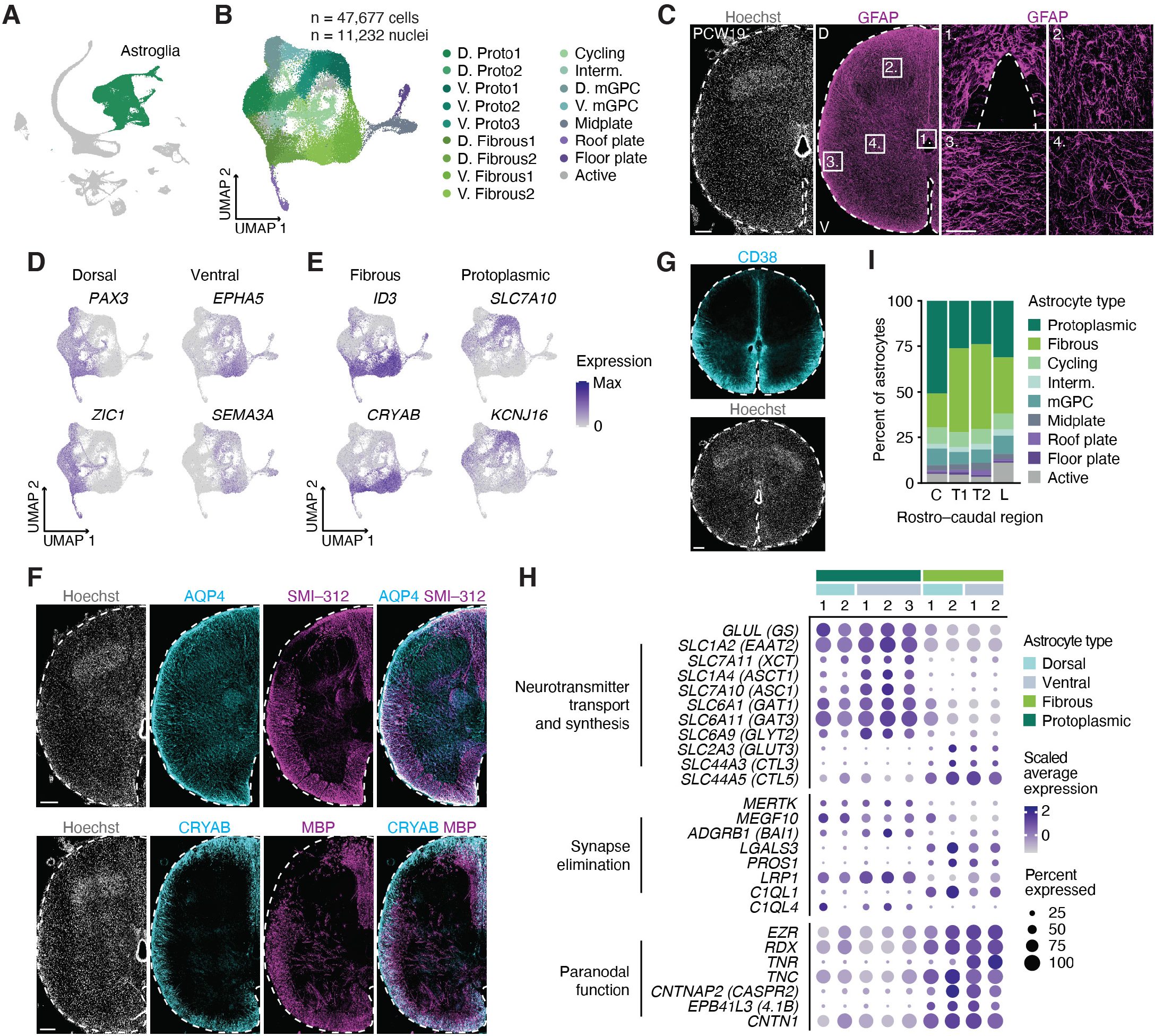
Astroglia in the spinal cord show diversity along several axes. **(A)** Highlight of astroglia in the main UMAP, showing the cells selected for subclustering. **(B)** UMAP of the astroglia subcluster colored by type. D, dorsal; V, ventral; Proto, protoplasmic; Interm., intermediate. **(C)** Representative immunohistochemistry image for the astroglia marker GFAP in a coronal upper thoracic spinal cord cryosection at PCW19. Insets show different types of astroglia in the spinal cord. **(D)** UMAP plots showing gene expression of markers associated with dorsal and ventral identities in the astroglia subcluster. **(E)** UMAP plots showing gene expression of markers associated with fibrous and protoplasmic astrocyte subtypes. **(F)** Representative immunohistochemistry images showing the general astrocyte marker AQP4 and the fibrous astrocyte marker CRYAB in the white matter of the spinal cord with neuronal axons expressing SMI-312 and oligodendrocytes expressing MBP. **(G)** Representative immunohistochemistry image showing CD38 in a coronal spinal cord cryosection at PCW19. **(H)** Dot plot showing the expression of selected genes associated with astrocyte functions in fibrous and protoplasmic subtypes. The size of the dots represents the percent of cells expressing each gene while the color depicts the scaled average expression per subtype. **(I)** Bar plot showing the proportion of each astroglia subtype in cervical (C), thoracic (T1, T2) and lumbar (L) regions in PCW18 spinal cord. Scale bars: 50 μm (insets in **C**), 200 μm (**C**, **F**, **G**).

The first axis corresponded to dorso-ventral positioning, with some astroglia expressing dorsal-related genes (*PAX3, ZIC1*), and others expressing ventral-related genes (*EPHA5, SEMA3A*; **Fig. 3D**), in line with what was described in the mouse spinal cord (Molofsky et al., 2014). Within the ventral group, we also found a population of V1 astrocytes characterized by the expression of *RELN* and the absence of *SLIT1* (Hochstim et al., 2008) (**Fig. S5A–C**). We validated the dorso-ventral division by immunohistochemistry with PAX3 and NKX6-1 (**Fig. S5D, E**). In addition, we found that homeodomain and forkhead transcription factors that are generally associated with positional identity in the spinal cord were expressed in a subtype-specific pattern (e.g. *PAX7, IRX2, DBX2, FOXP2, NKX6-1, NKX2-2* and *NKX6-2*) (**Fig. S5F**). The second axis of diversity corresponded to white matter and grey matter astrocytes (Oberheim et al., 2009) (**Fig. S6A–C**; **Table S9–11**). White matter or fibrous astrocytes were characterized by the expression of *CRYAB* and *ID3*, as well as higher expression of *GFAP* (Bakken et al., 2021; Hodge et al., 2019; Oberheim et al., 2009) (**Fig. 3E**, **S6A**). We validated the presence of these astrocytes in the white matter and found that CRYAB is specifically expressed in regions where myelinated SMI-312 axons are present (**Fig. 3F**). Grey matter or protoplasmic astrocytes were characterized by the expression of the amino acid transporter *SLC7A10* and the inward-rectifying potassium channel *KCNJ16* (*Kir5.1*; **Fig. 3E**). Further examination of these axes of division allowed us to discover new astrocyte markers. For example, we found that the transmembrane glycoprotein CD38 was specifically expressed in ventral fibrous astrocytes (**Fig. 3G, S6D**).

Astrocytes play crucial roles in the development and maintenance of neuronal function including neurotransmitter and potassium homeostasis, synapse formation and elimination, and blood brain barrier (BBB) function (Allen, 2014; Allen and Barres, 2009; Allen and Eroglu, 2017; Allen et al., 2012; Chung et al., 2013). Therefore, we next plotted genes associated with astrocyte function and asked if astrocyte positional or anatomical diversity was linked to functional diversity (**Fig. 3H**, **S6E**). We found that all types of astrocytes express Na^+^-K^+^-ATPases and potassium channels; however, different astrocyte subtypes specifically expressed different types (e.g. fibrous: *ATP1A2, ATP1B2, ATP1B1* and *ATP1A1*; protoplasmic: higher *ATP1A2* and *ATP1B2*). We also found that different astrocyte subtypes express different ionotropic and metabotropic receptors (fibrous: *GRM7* and *GRM8*; protoplasmic: *GRM3* and *GRM5*). Moreover, while most astrocytes are highly permeable to Ca^2+^, dorsal protoplasmic astrocytes are not, since they specifically express the AMPA receptor subunit GRIA2 (*GluA2*) (Droste et al., 2017; Man, 2011). Finally, we found that protoplasmic astrocytes have a higher expression of neurotransmitter transport genes (e.g. *SLC1A2*, *SLC7A10*, *SLC6A9*) suggesting a more prominent role in synaptically-released neurotransmitter clearance. In contrast, we found that fibrous astrocytes expressed genes involved in the regulation of neuronal transmission at the node of Ranvier. For example, they highly express the membrane proteins ezrin (*EZR*) and radixin (*RDX*), which mediate the motility of peripheral astrocyte processes (Lavialle et al., 2011). We also found both secreted extracellular matrix-binding proteins tenascin-C (*TNC*) and tenascin-R (*TNR*) were highly expressed in fibrous astrocytes. TNC and TNR bind and cluster sodium channels at the nodes of Ranvier (Srinivasan et al., 1998; Xiao et al., 1999). Based on this data and on their physical position in the white matter, we next wondered if fibrous astrocytes might be interacting with oligodendrocytes to play other roles in the spinal cord. We performed a cell-cell interaction analysis using the tool NATMI and found that ventral fibrous astrocytes and MOL showed the highest specificity of interactions (**Fig. S6F, G**; **Table S12**), which suggested that crosstalk between these two cell types might play roles in trophic support, cell adhesion and guidance, and BBB homeostasis.

Finally, we wondered if astrocyte diversity was similar along the rostro-caudal axis. We used data from one sample that was split into cervical, thoracic and lumbar regions (**Fig. S1D–F**). Interestingly, we found that although most astroglia subtypes were distributed equally within these four regions, protoplasmic astrocytes were enriched in the cervical region (**Fig. 3I**), and this difference was in part due to the presence of the V. Proto2 cluster almost exclusively in the cervical region, indicating that astrocyte diversity is present along multiple axes in the spinal cord (**Fig. S6H, I**).

### Midline glia and mGPCs in the developing human spinal cord

We next explored the diversity of progenitors. First, we focused on cells lining the ventricular zone (VZ), which included roof plate (RP), floor plate (FP) and midplate (MP) cells (**Fig. 4A, B**; **Table S13**). RP cells expressed *GDF7, GDF10, ZIC1* and *MSX1* (**Fig. 4C, S7A**), and FP cells expressed *SHH* and *FOXA2* (**Fig. 4C**) (Ghazale et al., 2019), as well as PAX7 as recently described (Rayon et al., 2021) (**Fig. S7B**). We validated ZIC1, FOXA2 and PAX7 expression in cryosections (**Fig. S7C, D**). Surprisingly, we found FOXA2^+^ FP cells dissociating from the VZ wall (**Fig. S7D**), a process that in mice takes place starting at E16.5 (Cañizares et al., 2020). We also noticed that although cells in the VZ showed distinct domains of expression along the dorso-ventral axis (**Fig. S7E–H**), this organization was different to that present at earlier stages (Rayon et al., 2021; Sagner and Briscoe, 2019). For example, we found that the p3, pMN and p2 marker NKX6-1 was not restricted to ventral domains, but was instead expressed along the entire MP and FP (**Fig. S7G, H**). Moreover, the pMN marker OLIG2 was not expressed in the VZ at this stage (**Fig. S7I, J**), although oligodendrogenesis is still ongoing (Weidenheim et al., 1994).

**Figure 4:**
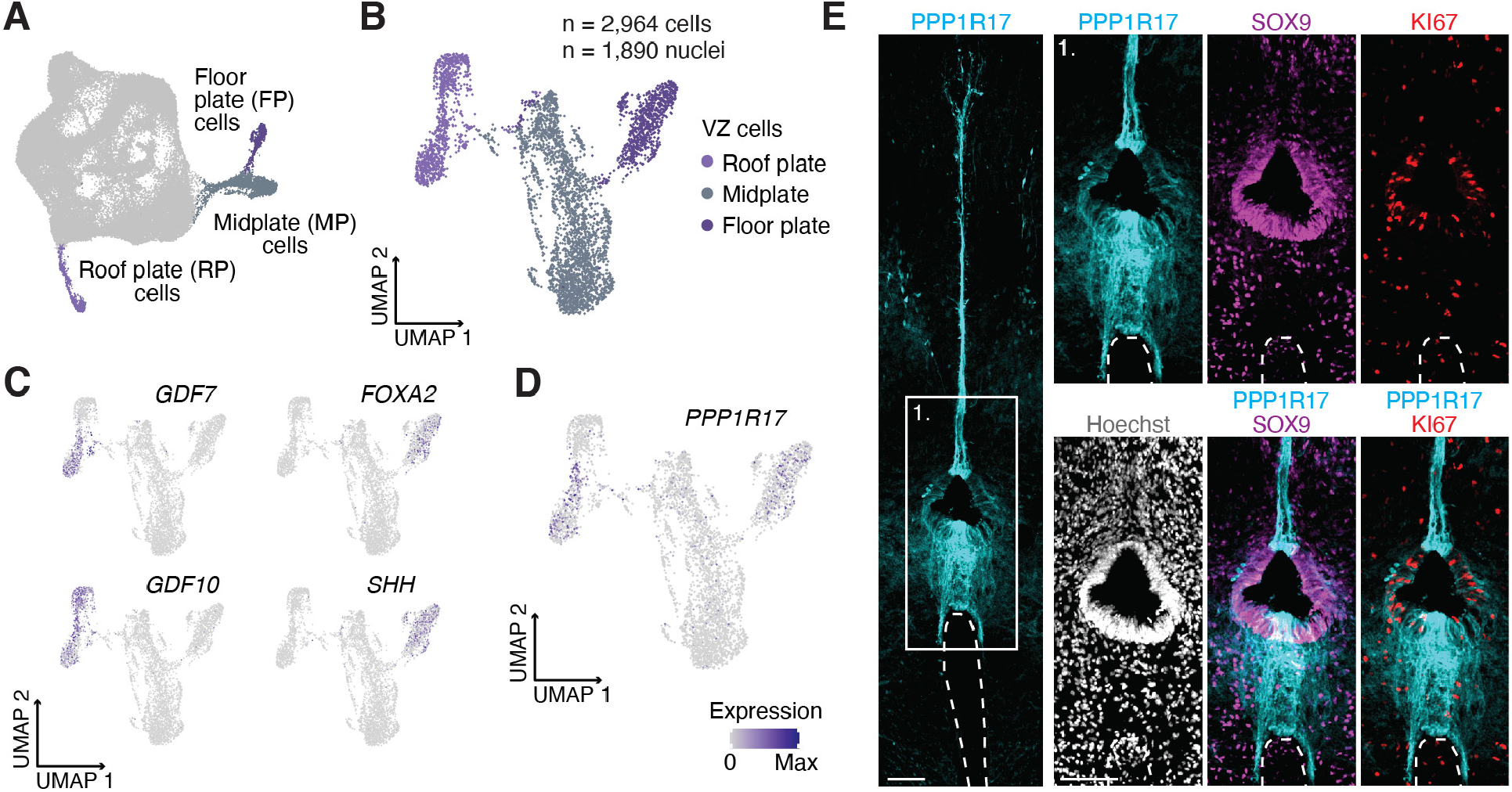
Midline glia in the developing human spinal cord. **(A)** Highlight of ventricular zone (VZ) cells in the astroglia subcluster, highlighting the cells selected for subclustering. **(B)** UMAP of the VZ cell subcluster, colored by cell type. **(C)** UMAP plots showing gene expression of markers specific to roof plate (RP; *GDF7, GDF10*) and floor plate (FP; *FOXA2, SHH*) cells. **(D)** UMAP plot showing expression of *PPP1R17* in RP and FP cells. **(E)** Representative immunohistochemistry images showing PPP1R17, SOX9 and the cell cycle marker KI67 in VZ cells in a coronal upper thoracic spinal cord cryosection. Hoechst shows nuclei. Scale bars: 100 μm (**E**).

Closer inspection of the FP and RP cells revealed expression of the phosphatase inhibitor *PPP1R17* (**Fig. 4D**), a marker for intermediate progenitor cells in the cerebral cortex (Trevino et al., 2021) with human-specific gene regulation (Girskis et al., 2021). We performed immunohistochemistry and were surprised to find *PPP1R17* expressed in a group of FP and RP cells and along the dorsal and ventral midline (**Fig. 4E**). Their position and radial-like morphology extending from the VZ to the pia is suggestive of a midline glia-like or Nestin^+^ radial glia identity (Ghazale et al., 2019; Shinozuka and Takada, 2021). Midline glia in the Drosophila embryo express highly conserved guidance molecules and play important roles in directing axonal crossing (Lemke, 2001). Analysis of FP and RP *PPP1R17*^+^ cells indicated that they express ligands and receptors important for midline crossing (Chédotal, 2019), such as Netrin (*NTN1*), SLITs, semaphorins and ephrins (eg. *EFNB3*; **Fig. S7K**; **Table S14, S15**). Moreover, immunohistochemistry with the axonal marker SMI-312 and the sensory neuron marker substance P showed axons crossing both the dorsal and ventral midlines at PCW19 (**Fig. S7L, M**).

In the midgestation human cerebral cortex, we identified a group of multipotent EGFR^+^/OLIG2^+^/ASCL1^+^ glial progenitors (mGPC) that may generate both astrocytes and oligodendrocytes (Trevino et al., 2021). Here, we found a similar population expressing *EGFR, OLIG2* and *ASCL1* (**Fig. S8A, B**) that formed a bridge between the astroglia and OPC/Oligo clusters, suggestive of multipotency; half of these cells expressed astroglia markers (*SOX9*, *AQP4*) while the other half expressed OPC/Oligo markers (*SOX10, PDGFRA*) (**Fig. S8C**). Immunohistochemistry for EGFR and OLIG2 identified cells positioned mainly in the intermediate grey matter adjacent to the VZ (**Fig. S8D, E,** inset 1) expressing either the astroglia marker SOX9 or the oligodendrocyte lineage marker NKX2-2, consistent with mGPCs.

### Motor neuron identity is specified at midgestation in human

Next, we investigated cell diversity in spinal cord neurons (**Fig. 5A**). We subclustered all neurons and found groups with different neurotransmitter identities (**Fig. 5B–D, S9A– D**; **Table S16**), including glutamatergic (Glut: *SLC17A6*), GABAergic and glycinergic (GABA and GABA/Glyc.: *GAD1*, *GAD2*, *SLC32A1*, and *SLC6A5*), and cholinergic (*SLC5A7*, *SLC18A3* and *CHAT*). We validated the presence of these neuronal types by immunohistochemistry (**Fig. 5E**). Next, we used label transfer to compare our cell types to mouse datasets (Häring et al., 2018; Osseward et al., 2021; Rosenberg et al., 2018; Sathyamurthy et al., 2018; Zeisel et al., 2018). This revealed the presence of different types of excitatory, inhibitory and cholinergic neurons (**Fig. 5F, S9E, F**; **Table S17**), as well as dorsal, ventral and intermediate types (**Fig. 5G, S9G, H**).

**Figure 5:**
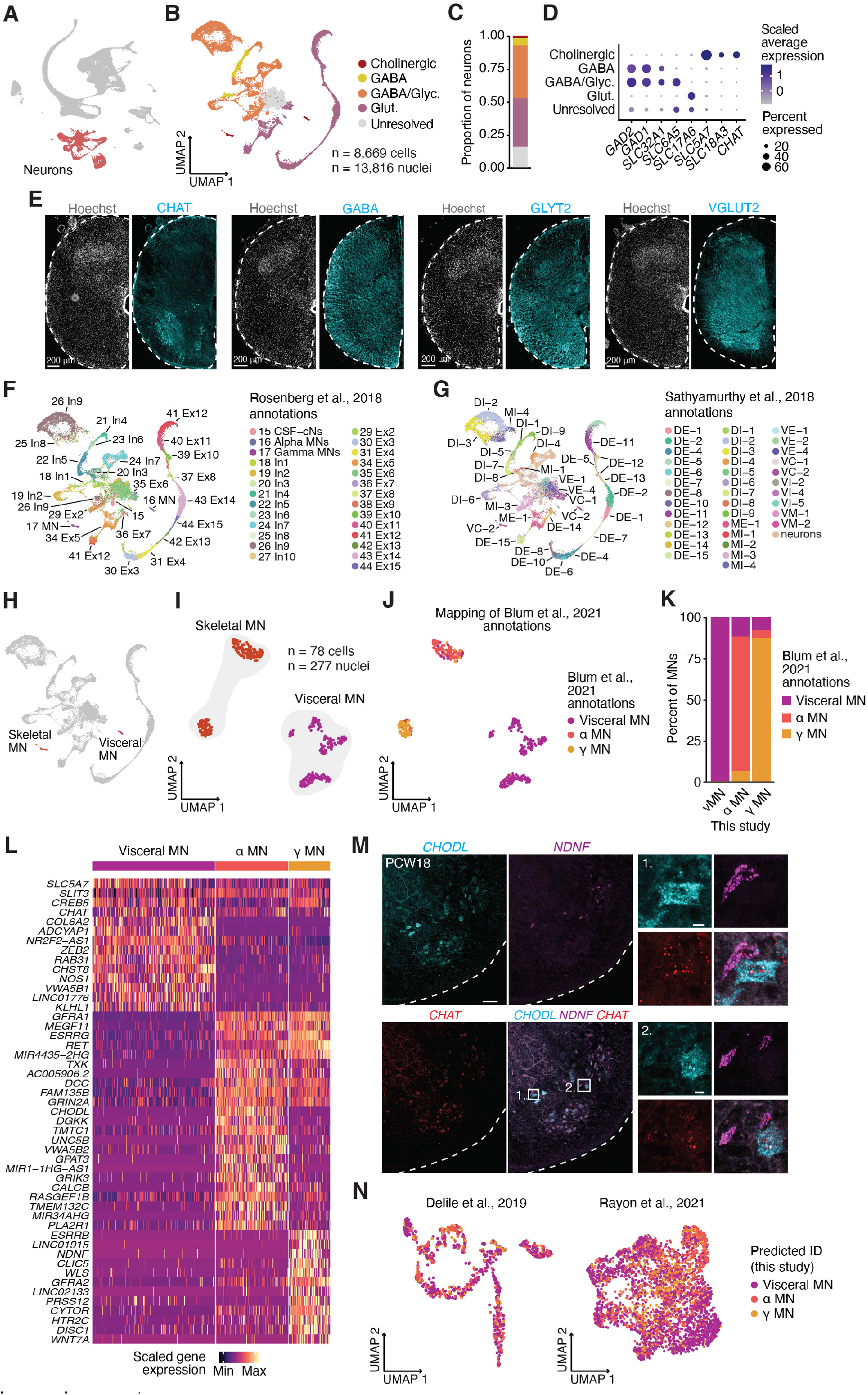
Neuron and motor neuron diversity in the spinal cord. **(A)** Highlight of neurons in the main UMAP, highlighting the cells selected for subclustering. **(B)** UMAP of the neuron subcluster, colored by neurotransmitter identity. **(C)** Bar plot showing the proportion of cells within the neuron subcluster in each neurotransmitter identity group. **(D)** Bubble plot showing expression of neurotransmitter identity-related genes as average scaled expression per group. *SLC32A1, SLC6A5, SLC17A6, SLC5A7* and *SLC18A3* are also known as *VGAT, GLYT2, VGLUT2, CHT1* and *VACHT*, respectively. **(E)** Representative immunohistochemistry images showing neurons with different neurotransmitter identities (CHAT shows cholinergic neurons, GLYT2 shows glycinergic. neurons, GABA shows GABAergic neurons and VGLUT2 also known as SLC17A6 shows glutamatergic neurons) in coronal spinal cord cryosections at PCW19. **(F)** Label transfer showing neuronal annotations from Rosenberg et al., 2018 in the neuron subcluster. **(G)** Label transfer showing neuronal annotations from Sathyamurthy et al., 2018 in the neuron subcluster. **(H)** Highlight of skeletal and visceral motor neurons (MN) in the neuron subcluster. **(I)** Motor neuron subcluster colored by identity. **(J)** Motor neuron subcluster showing the predicted identities of cells based on Blum et al., 2021 label transfer. **(K)** Bar plot showing the percent of predicted identities in the motor neuron subcluster (this study) based on annotations from Blum et al., 2021. **(L)** Heatmap of scaled gene expression across cells in the MN subcluster. The parameter min.diff.pct in ‘FindMarkers’ was set to 0.3 to select genes specific to each group. **(M)** Representative *in situ* hybridization of *CHAT, CHODL* and *NDNF* in a coronal lumbar spinal cord cryosection at PCW18. Insets show *CHAT*^+^ motor neurons that express either the alpha marker *CHODL* or the gamma marker *NDNF*. **(N)** Label transfer showing predicted IDs from motor neuron clusters in this study onto Delile et al., 2019 motor neurons (left) and Rayon et al., 2021 motor neurons (right). Scale bars: 10 μm (insets in **M**), 100 μm (**M**).

We next focused on cholinergic neuron diversity. Of the two cholinergic clusters we identified, one expressed visceral motor neuron (MN) markers (*NOS1*) and the other expressed markers of skeletal MNs (*RET* and *GFRA1;* **Fig. 5H**, **S10A; Table S18**). Subclustering revealed that skeletal MNs were composed of two subgroups (**Fig. 5I**). Using label transfer of MNs from adult mouse spinal cord (Blum et al., 2021), we found that PCW17-18 visceral MNs mapped onto mouse adult visceral MNs, while skeletal MNs were divided into α and γ subgroups (**Fig. 5J–L, S10B–G**; **Table S19**, **S20**). Comparison of human α and γ MNs with their adult mouse counterparts revealed some similarities (**Fig. S11A, B**, **Table S21**, **S22**). For example, αMNs expressed *CHODL, SV2B* and *VIPR2* in both datasets, while γMNs expressed *ESRRB* and *WLS*. In addition, we found differences among the two datasets that, in part, might highlight the different ages sampled. For example, we found that unlike adult mouse αMNs, human PCW17-18 αMNs expressed several type II cadherins (*CDH7, CDH9, CDH10*, *CDH20*) that might regulate motor pool segregation (Dewitz et al., 2019; Price et al., 2002; Vagnozzi et al., 2020). Other differences included the expression of *UNC5B, TXK* and *DGKK* in human αMNs, and the expression of *NDNF*, *CLIC5* and *GFRA2* in human γMNs. To validate the expression of some of these markers in human MNs, we performed RNAScope for the α marker *CHODL* and the novel γ marker *NDNF* at PCW18 and PCW19. We found that *CHODL* and *NDNF* were expressed in *CHAT*^+^ cells in the ventral horn, but did not colocalize (**Fig. 5M**, **S11C**), indicating their presence in different types of MNs. Interestingly, we did not see this separation between *Chodl* and *Ndnf* in mouse spinal cord sections at P0 or P25 (**Fig. S11D, E**). At both ages *Chodl* was expressed in a proportion of MNs, but *Ndnf* was either not expressed or expressed at low levels in *Chodl* neurons, indicating that *NDNF* is a human-specific marker for γMNs.

MN diversification into extrafusal- and intrafusal-innervating αMNs and γMNs is only apparent at late stages of rodent development, with the earliest marker of γMNs detectable at E17.5 (Ashrafi et al., 2012), so we were surprised to see this separation at early midgestation in human. To determine whether the absence of early markers for αMNs and γMNs hindered identification of these populations at earlier time points, we used α, γ and visceral MN markers identified in this study and performed label transfer onto mouse and human datasets at earlier stages (E9.5–E13.5, Delile et al., 2019; and CS12–CS19 or PCW4–PCW7, Rayon et al., 2021). Interestingly, we did not see clear separations between these three MN groups at early developmental stages (**Fig. 5N, S12A–H**).

### Mapping disease-related genes

We next mapped the expression of genes associated with disease in our spinal cord dataset. First, we focused on genes associated with myelin-related disorders, including leukodystrophies and degenerative white matter disorders (**Fig. S13A, Table S23**). We found a subgroup of genes expressed specifically in myelinating oligodendrocytes or Schwann cells (*PLP1*, *FA2H* and *GJC2*) and a subgroup expressed in different types of astrocytes (*GJA1, GFAP* and *EDNRB*), or in microglia (*CSF1R* and *TREM2*), suggesting that interactions between different cell types may contribute to pathogenesis (van der Knaap and Bugiani, 2017). Next, we focused on genes linked to the Charcot-Marie-Tooth neuropathy (CMT, **Fig. S13B**). We found a group of genes expressed in the OPC/Oligo lineage or Schwann cells (*MPZ* and *PMP22*), astrocytes (*FGD4*) and microglia (*EGR2*), as well as genes that appeared to be ubiquitously expressed (*KIF1B* and *SBF2*). Lastly, we mapped the expression of genes associated with ALS, which is characterized by the selective loss of motor neurons (**Fig. S13C**). We found that, although some genes appeared to be highly expressed in neurons (*MAPT* and *UNC13A*) or specifically expressed in motor neurons (*NEFH* and *PRPH*), the majority of them were ubiquitously expressed (*SOD1*), or present in non-neuronal cell types such as OPC/Oligo (*MOBP*), or VLMC, pericytes and microglia (*SQSTM1* and *GRN*), highlighting the role of non-cell autonomous toxicity in this disease (Ilieva et al., 2009).

## Discussion

Single cell transcriptomics technologies have enabled the systematic profiling of cell diversity across the central nervous system at an unprecedented rate (Winnubst and Arber, 2021). To generate a comprehensive census of cell types of the midgestation human spinal cord, we performed single cell and single nucleus RNA sequencing.

A common theme that emerged through our analysis was the existence of a transcriptional positional identity code governing cell diversity. Our data suggest that the acquisition of positional identity is a common fundamental principle guiding patterning in the spinal cord beyond neuronal specification. Astrocytes displayed the most striking example of this, but cells in the VZ, mGPCs and OPCs also showed position-related signatures. In astrocytes, positional differences appeared to be linked to functional differences based on gene expression; however, further work is needed to demonstrate this. Sampling at earlier timepoints could also determine the relationship between some of the cell types we described. For example, how fibrous and protoplasmic astrocytes are generated during development or if they are derived from a common progenitor remains unknown. Similarly, the dual lineage potential of mGPCs is reminiscent of ependymal cells in the adult murine spinal cord that upon injury are activated to generate both oligodendrocytes and astrocytes (Llorens-Bobadilla et al., 2020), yet whether mGPCs are generated from VZ/ependymal cells during development is unclear.

Previous reports suggested that the human spinal cord develops and matures faster when compared to rodents (Ohmura and Kuniyoshi, 2017; Tadros et al., 2015). Our analysis indeed revealed that processes that happen towards the end of development in mice take place at midgestation in human. First, we found that gliogenesis and ventricular layer remodeling–processes that start in the last phase of uterine development in mice (Cañizares et al., 2020; Tait et al., 2020; Zhou and Anderson, 2002), are in place at PCW17-18 in human. In addition, we discovered that diversification of MNs into α and γ takes place in early development (by PCW17), unlike in mice (Ashrafi et al., 2012). This acceleration of developmental processes in the human spinal cord is in contrast to the slower pace observed in the human cerebral cortex (Somel et al., 2009; Sousa et al., 2017). Evidence for early muscle innervation by motor neurons in human compared to rodents may explain, in part, this discrepancy (Tadros et al., 2015). A remaining challenge will be to comprehensively compare developmental landmarks and cellular diversity across species to identify unique features and uncover the molecular machinery underlying maturation at a different developmental pace.

## Acknowledgements

We thank members of S.P. Pașca, A.M. Pașca and W.J. Greenleaf laboratories for support, discussion and advice, especially S. Kanton, Y. Miura, L. Li, A. Trevino, K.W. Kelley, M. Onesto and F. Birey. This work was supported by the S. Coates and V. Coates Foundation (S.P.P.), the Stanford Brain Organogenesis Program and the Big Idea Grant in the Stanford Wu Tsai Neurosciences Institute (S.P.P.), Bio-X (S.P.P.), the Kwan Fund (S.P.P), the Senkut Research Funds (S.P.P.), the Chan Zuckerberg Initiative Ben Barres Investigator Award (S.P.P), the Stanford University Department of Neurosurgery (J.A.K.) and the Stanford Wu Tsai Neurosciences Institute (J.A.K.). S.P.P. is a New York Stem Cell Foundation Robertson Stem Cell investigator. W.J.G. is a Chan Zuckerberg Biohub investigator and acknowledges grants 2017-174468 and 2018-182817 from the Chan Zuckerberg Initiative. Fellowship support was provided by the Idun Berry Postdoctoral Fellowship (J.A.).

## Author contributions

J.A., N.T. and S.P.P. conceived the project and designed experiments. N.T. performed data analysis with guidance from F.M. and W.J.G. J.A. guided the biological interpretation of the analysis and performed immunohistochemistry validations. F.M. performed the RNA velocity analysis. J.L.S and J.A.K. performed and interpreted RNAScope validations. A.M.P., N.D.A. and J.A. processed samples. X.C. performed the nuclei NeuN-sort experiment. S.J.Y. performed Thy1 immunopanning. J.A. performed Chromium 10x. J.A., N.T. and S.P.P. wrote the manuscript with input from all authors. J.A. and S.P.P. supervised the work.

## Declaration of interests

W.J.G. was a consultant for 10x Genomics.

## Methods

### Data and materials availability

Data used for the analyses presented in this work are available under GEO accession number GSE188516. A website associated with the manuscript, including an interactive data browser, is available at https://devspinalcord.su.domains/.

### Human tissue

De-identified spinal cord samples were obtained at Stanford University School of Medicine from elective pregnancy terminations under a protocol approved by the Research Compliance Office at Stanford University. Samples were delivered on ice and processed for single cell analyses or immunocytochemistry within three hours of the procedure.

### Sample collection and single cell data generation

Single cell dissociation and Thy1 immunopanning was performed as previously described (Sloan et al., 2018; Trevino et al., 2020, 2021). Briefly, spinal cords were dissected out of the vertebral column, chopped and incubated in 30 U/mL of papain enzyme solution (Worthington, LS03126) for 45 min at 37°C. After digestion samples were washed with a protease inhibitor solution and gently triturated using progressively smaller pipette tips to achieve a single cell suspension. Cells were resuspended in 0.02% BSA/PBS and passed through a 70 μm flowmi filter and either continued to single cell sample preparation or Thy1 immunopanning to enrich for neurons. For immunopanning, the single cell suspension was added to a plastic petri dish pre-coated with an anti-Thy1 antibody (CD90, BD Biosciences, 550402) and incubated for 10-30 minutes at room temperature. Bound cells were incubated in an Accutase solution (Innovative Cell Technologies, AT104) at 37°C for 3-5 minutes, and then gently washed off, spun down at 200 x g for 5 minutes and resuspended in cold 0.02% BSA/PBS.

Nuclei isolation was performed as described in (Matson et al., 2018) with some modifications. Briefly, dissected spinal cords were disrupted using the detergent-mechanical cell lysis method using a 2 ml glass tissue grinder (Sigma-Aldrich/KIMBLE, D8938). Crude nuclei were then filtered using a 40 μm filter and centrifuged at 320 x g for 10 minutes at 4°C before performing a sucrose density gradient to separate them from cellular debris. After a centrifugation step (320 x g, 20 minutes at 4°C), samples were resuspended in 0.04% BSA/PBS supplemented with 0.2 U/μl RNAse inhibitor (Ambion 40U/μl, AM2682) and passed through a 40 μm flowmi filter. NeuN-sorting was performed as described in (Matevossian and Akbarian, 2008). Briefly, the cell suspension was mixed with 1.2 μl of mouse anti-NeuN antibody (Millipore, MAB377) and 1 μl of anti-mouse Alexa Fluor 488 (Thermo Fisher Scientific, A21202) in PBS with 0.5% BSA and 10% normal donkey serum for 45 minutes on ice. The sorting was performed on a BD FACS Aria II Cell Sorter at the Stanford Shared FACS Facility. A 100 μm nozzle and Purity mode were used during the sorting. Nuclei suspension stained with secondary antibody only was used as a control for setting up the gate. Sorted nuclei were collected in 0.04% BSA/PBS supplemented with 0.2 U/μl RNAse inhibitor.

Cell and nucleus suspensions were then loaded onto a Chromium Single cell 3’ chip, v3 (10x Genomics) and processed according to instructions with a target of 10,000 cells, 8-10,000 nuclei and 4,000 NeuN-sorted nuclei.

### Analysis and processing of scRNA and snRNA sequencing data

Single cell (sc) and single nucleus (sn) libraries were sequenced by Admera Health on a Novaseq S4 (Illumina) using 150 x 2 chemistry. Fastq files were obtained from Admera Health and subsequently aligned to human reference genome GRCh38 (2020-A) using Cell Ranger (v5.0.0, 10x Genomics) on the Sherlock Stanford Computing Cluster. We used the ‘include introns’ option for both single cell and single nucleus samples and all other parameters were kept as default. Spliced and unspliced counts were computed using velocyto [v0.17.17, (La Manno et al., 2018)] from the BAM files produced by Cell Ranger with default parameters and the same reference used for alignment.

Analysis of the counts-by-gene matrix created by Cell Ranger was performed using ‘Seurat’ [v3.9.9.9024, beta of v4; (Stuart et al., 2019)] in R (v3.6.1). Quality control (QC) and filtering was performed on single cells and single nuclei separately. These were first filtered to remove cells or nuclei with fewer than 1,000 features or greater than 30% mitochondrial reads. Subsequently, through repeated rounds of clustering and QC, we removed doublets and low-quality cells. For each round of clustering and QC, known cell type-specific marker genes were used to annotate cells. We used *SOX10, OLIG2, PDGFRA, MOG* and *MBP* to identify cells in the OPC/Oligo lineage; *AQP4, GFAP, FGFR3* and *SOX9* to identify astroglia; *STMN2* and *MYT1L* to identify neurons; *AIF1* for microglia; *TOP2A* for cycling cells; *PECAM1* for endothelial cells; *NGFR* for Schwann cells; and *COL1A1* for VLMC. Clusters containing a majority of cells that expressed well known cell-type specific markers for two or more cell types (e.g. clusters expressing both *MYT1L* and *SOX10* or *AQP4, FGFR3* and *AIF1*) were assumed to be doublets and removed. We also removed clusters that were low-quality, which was determined by a majority of cells with low nCount and high mitochondrial percent. From the 138,159 single cells identified by Cell Ranger, 25,605 were identified as doublets or low-quality cells and removed. Similarly, 9,247 nuclei were removed out of the 44,131 initially identified by Cell Ranger.

Remaining single cells were then normalized, scaled, and integrated to correct for batch effects using the Integrate pipeline from Seurat, with 3,000 integration features used across the two batches of cells we collected. Single nuclei were also normalized, scaled, and integrated with 2000 integration features. PCA analysis was performed with the first 75 and first 50 principal components selected for single cells and nuclei, respectively. These principal components were then used in the ‘FindNeighbors’ and ‘FindClusters’ functions to determine cell groupings and to generate a two-dimensional UMAP projection via ‘FindUMAP’ for each dataset. These functions were run with the parameters recommended in Seurat’s tutorial for large datasets (‘FindNeighbors’ nn.eps = 0.5) and were parallelized using 4 threads using the R package ‘future.’

After QC and filtering we then integrated the two datasets together to generate our final UMAP. To aid the integration, we removed mitochondrial and ribosomal genes from the count matrices before integration. We used 2,000 features for CCA integration and 60 dimensions along with parameters in the pipeline described above for ‘FindNeighbors.’ This produced a total of 53 clusters which were then combined and manually annotated based on the cell-type specific markers described above. The astrocyte and oligodendrocyte subclusters were generated by rerunning ‘FindVariableFeatures’, ‘ScaleData’, ‘RunPCA’, ‘FindNeighbors’, ‘FindClusters’, and ‘RunUMAP’ on the integrated assay of the combined dataset. The mGPC cluster was split between the astroglia and OPC/Oligo subclusters: Seurat cluster 7 was included with the astroglia and Seurat cluster 46 with the OPC/Oligo. For neurons we performed a further round of doublet removal and then reintegrated the data due to the prevalence of Thy1 immunopanned cells and NeuN-sorted nuclei, integrating across collection type instead of sequencing run. Within the astroglia and neuron subclusters, we further subclustered cycling astroglia (10 dims, 1000 variable features) and motor neurons (8 dimensions, 750 features) respectively. In the motor neuron subcluster we removed a cluster of cells (n = 15) that expressed DRG markers. Annotation of subgroups or cell types within subclusters was determined by gene expression of known markers or label transfer with published datasets. In some cases, several Seurat clusters were combined to form a subgroup, and the identity of these is shown in Supplementary Tables.

For heatmaps showing cluster expression we pseudobulked each cluster by summing up gene expression before normalization via Seurat’s ‘NormalizeData.’ Cell cycle scores were generated in Seurat using the ‘CellCycleScoring’ function and the default ‘s.genes’ and ‘g2.genes’ objects loaded in Seurat. For MA plots, differential expression was computed with ‘FindMarkers’ with the option ‘logfc.threshold’ set to 0.

Myelin-related disorder genes were selected from: (Abrams and Scherer, 2012; Brenner et al., 2001; Charzewska et al., 2016; Depienne et al., 2013; van der Knaap and Bugiani, 2017; Zardadi et al., 2021). Charcot-Marie-Tooth (CMT)-associated genes were gathered from the Human Phenotype Ontology website (https://hpo.jax.org/app/). Amyotrophic lateral sclerosis-associated genes were selected from human exome sequencing studies: (Cirulli et al., 2015; Farhan et al., 2019; Kenna et al., 2016; Nicolas et al., 2018; van Rheenen et al., 2016; Smith et al., 2014, 2017).

### Differential expression and GO analysis

Differential Expression was performed in Seurat with either ‘FindAllMarkers’ or ‘FindMarkers’ using default parameters except that ‘min.pct’ was changed to 0.25, unless otherwise specified. All tests were done using the Wilcoxon rank sum test. Top markers were selected from those that had *P*adj values < 0.01, then ranked by their log fold change.

GO analysis for differential expression between single cells and single nuclei was completed using ‘fgsea’ (v1.19.2) with terms supplied via the GSEA mSigDB, specifically those related to Biological Processes (BP). To find terms that were enriched, an overexpression analysis was carried out with a hypergeometric test using the ‘fora’ function. The gene universe was taken to be all genes present in the RNA counts matrix, and the gene list was taken to be at most the top 250 genes by log fold change with adjusted *P*-values < 0.01 based on results from differential expression. Calculated adjusted *P*-values were then used in subsequent analyses.

### Label transfer with reference datasets

Data for label transfer analysis performed in this study (Zeisel et al., 2018, Rosenberg et al., 2018, Sathyamurthy et al., 2018, Delile et al., 2019, Häring et al., 2018, Blum et al., 2021, Rayon et al., 2021) was obtained from GEO. The provided annotations and counts by gene matrices were then processed in the same manner as our data before comparison. For mouse datasets, genes were converted from mouse MGI or ENSEMBL to HGNC symbols via a list of mouse to human homologs (Skene et al 2018). In the case of the Rayon et al, 2021 dataset, integration, normalization, and clustering were performed on the full dataset for annotation (2000 Variable Features, 2000 integration features, 40 dimensions, 0.3 resolution, 30% mitochondrial gene cutoff, 1000 nCount cutoff). Motor neurons for this dataset were then identified based on the expression of *ISL1, MNX1* and *CHAT* and subclustered (1000 Variable Features, 10 dimensions, 0.3 resolution). Label transfer was then carried out with all reference datasets using a CCA projection of the dataset on the first 25 dimensions using variable features from our dataset from the integrated assay between the neuron or motor neuron subcluster of our dataset and the RNA assay of the reference dataset.

### Cell-cell interaction analysis

We used NATMI to investigate ligand receptor interactions between cell types. Counts were taken from the RNA assay of an object, then normalized using ‘NormalizeData’ with default parameters and transformed into CPM/TPM as demonstrated by the NATMI instructions. The command line tool was then used with Python (3.6.1) to both extract and visualize the edges of the ligand receptor interaction network. We used 32 threads, a specificity cutoff of 0.05, and a detection cutoff of 0.1. This provided a list of edges with information on their expression, specificity, and detection (**Table S12**). For visualization we selected edges that were between astrocyte and OPC/Oligo subtypes but not between either astrocytes and astrocytes or OPC/Oligo and OPC/Oligo.

### RNA velocity and pseudotime

We computed RNA velocity using custom R scripts interfacing with the ‘scVelo’ toolkit (v.0.2.3) (Bergen et al., 2020) via the ‘reticulate’ R-Python interface. For this, we exported the Velocyto-derived spliced and unspliced counts along with Seurat-derived PC and UMAP representations of single cells as ‘AnnData’ objects. We filtered the dataset using the scVelo function ‘pp.filter_and_normalize’ (parameters: min_shared_counts = 10, n_top_genes = 2,000) and computed moments using ‘pp.moments’ (n_pcs = 30, n_neighbors = 30). We then used ‘tl.velocity’ with mode = ‘dynamical’ to compute cell velocities and ‘tt.velocity_graph’ to compute a velocity graph. Potential root and end point cells for the trajectory were computed using ‘tt.terminal_states’. ‘tt.velocity_pseudotime’ was applied to compute cell pseudotime scores. We then reimported the scVelo-derived gene and cell annotations (gene velocities and other scVelo-inferred model parameters, as well as cell pseudotime, velocity projections, root and end point probabilities) into the metadata of the R-based Seurat objects. Cell-type specific velocity genes were obtained using scVelo’s ‘rank_velocity_genes’ function.

To facilitate aggregate analysis along pseudotime, we obtained pseudobulk samples by sorting cells based on their pseudotime scores and merging bins of 100 cells. For merging gene expression levels, counts for all cells assigned to a pseudobulk sample were summed and the data was renormalized using counts per million normalization.

### Immunohistochemistry

Immunohistochemistry was performed as previously described (Sloan et al., 2018; Trevino et al., 2020, 2021). In brief, PCW18 and PCW19 spinal cords were dissected out of the vertebral column and fixed for 2-3 hours at 4°C with 4% paraformaldehyde (PFA, Electron Microscopy Sciences), washed with PBS and transferred to a 30% sucrose solution for 48-72 hours. Once the samples had sunk in this solution they were embedded in an OCT/30% sucrose solution (1:1) and snap frozen in dry ice. Cryosections were obtained using a Leica cryostat set at 20-30 μm and mounted on Superfrost Plus Micro slides (VWR, 48311-703). On the day of staining, sections were then blocked and permeabilized for 1 hour at RT in blocking solution containing 10% normal donkey serum, 0.3% Triton-X in PBS and then incubated with primary antibodies at 4°C overnight. The primary antibodies used are shown on **Table S24**. Primary antibodies were washed using PBS and sections were incubated with Alexa Fluor secondary antibodies (1:1,000, Life Technologies) for 1 hour at RT. Hoechst 33258 (Thermo Fisher Scientific, H3569) was used to visualize nuclei. Sections were mounted with glass coverslips using Aqua Polymount (Polysciences, 18606-5). Images were taken using a Leica SP8 confocal microscope or a Keyance fluorescence microscope and processed using ImageJ (Fiji). All immunohistochemistry validations were performed in PCW18 and PCW19 samples.

### *In situ* hybridization (RNAScope)

PCW18 (n=1) and PCW19 (n=1) samples were processed as described above. Spinal cords from P0 (n = 4) and P25 (n = 3) mice were isolated by hydraulic extrusion. Mouse spinal cords were isolated by hydraulic extrusion. Tissues were then fixed in 4% PFA either for two hours or overnight and cryopreserved in 30% sucrose. Lumbar spinal cords were embedded in OCT Compound and 20 μm transverse sections were cut on a Leica CM3050 S Cryostat. For RNAScope frozen cryosections were hydrated in PBS and pretreatment of sections was performed as follows: slides were baked at 60°C for 45 minutes, post-fixed in 4% PFA at RT for 1 hour, dehydrated in 50%, 70%, 100%, 100% ethanol (5 minutes each). Sections were then incubated in hydrogen peroxide for 10 minutes and antigen retrieval was performed in a vegetable steamer for 4 minutes. Finally, slides were baked a final time for 45 minutes. Sections were processed with the RNAScope Multiplex Fluorescent V2 Kit (ACD Biosciences) per the manufacturer’s guidelines. The following probes were used: Hs-CHAT-C2 (450671-C2), Hs-CHODL (506601), Hs-NDNF-C3 (495251-C3), Mm-Chat-C2 (408731-C2), Mm-Chodl (450211) and Mm-Ndnd-C3 (447471-C3). All imaging was done on a Leica SP8 confocal, maximum projections were generated using LASX software and further processed using ImageJ (Fiji).

## Supplementary Figures

**Supplementary Figure 1:**
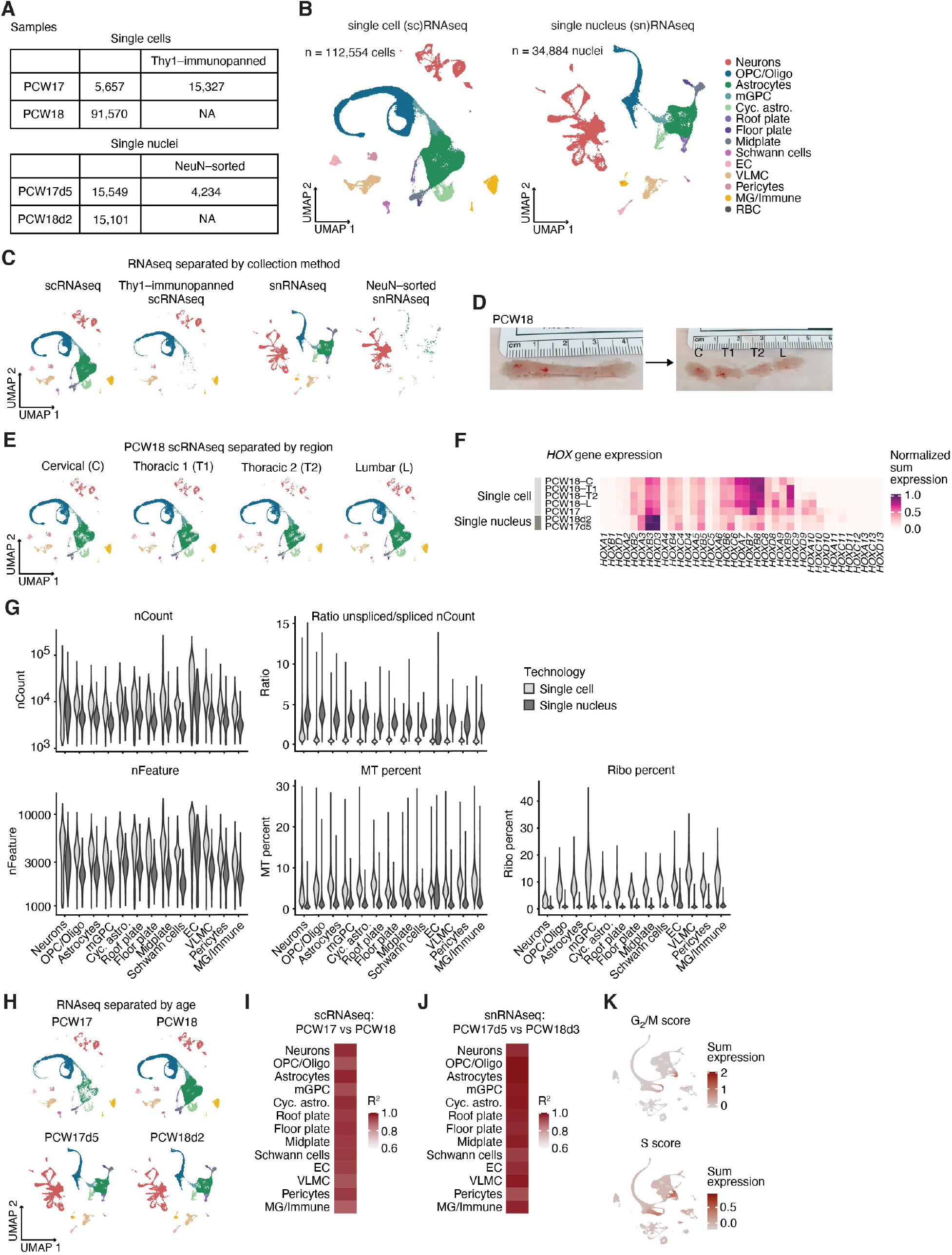
Sample overview and quality control metrics. **(A)** Table showing the number of single cells and single nuclei per sample and collection method. Numbers correspond to total numbers of cells or nuclei recovered after quality control (QC) and filtering. **(B)** UMAPs showing single cells (left) and single nuclei (right) before integration, colored by cell type. **(C)** Single cell and single nucleus UMAPs split by collection method. **(D)** Picture of PCW18 spinal cord sample before and after dividing it into rostro-caudal regions: cervical (C), thoracic (T1, T2) and lumbar (L). **(E)** Single cell UMAPs showing PCW18 cells split by region. **(F)** Heatmap showing *HOX* gene expression as the sum of gene counts normalized to total sample counts in all samples. **(G)** Violin plots showing total counts (nCount), ratio of spliced to unspliced counts, total number of features (nFeature), percent of mitochondrial (MT) counts, and percent of ribosomal (ribo) counts separated by cell type in both single cell and nucleus samples. **(H)** Single cell and single nucleus UMAPs split by sample age. **(I)** Heatmap showing the correlation of the normalized average gene expression between sample ages in single cell samples separated by cell type represented as R^2^ values. **(J)** Heatmap showing the correlation of the normalized average gene expression between ages in single nuclei samples separated by cell type represented as R^2^ values. **(K)** G2/M and S cell cycle scores in integrated single cell and single nuclei UMAP.

**Supplementary Figure 2:**
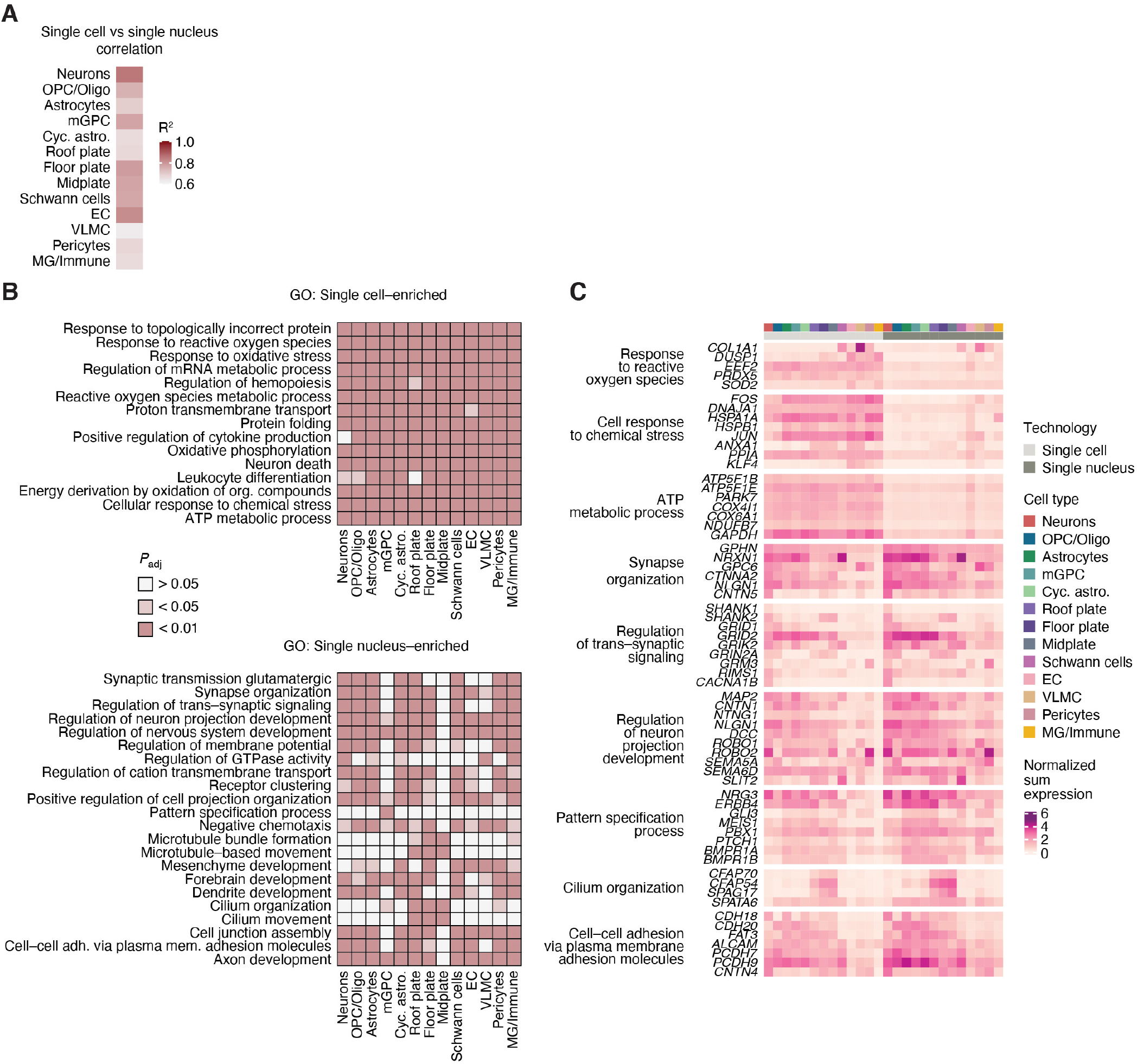
Comparison of single cell and single nucleus transcriptional data. **(A)** Heatmap showing the correlation of the normalized average gene expression between single cell and single nucleus samples separated by cell type represented as R^2^ values. **(B)** Heatmaps showing the top 5 gene ontology (GO) terms by cell type of genes enriched in either single cell samples (top) or single nucleus samples (bottom) per cell type, colored by significance level. **(C)** Heatmap showing gene expression of selected genes within GO terms that are differentially expressed in single cell and single nucleus samples as the sum of gene counts normalized to total counts per cell type.

**Supplementary Figure 3:**
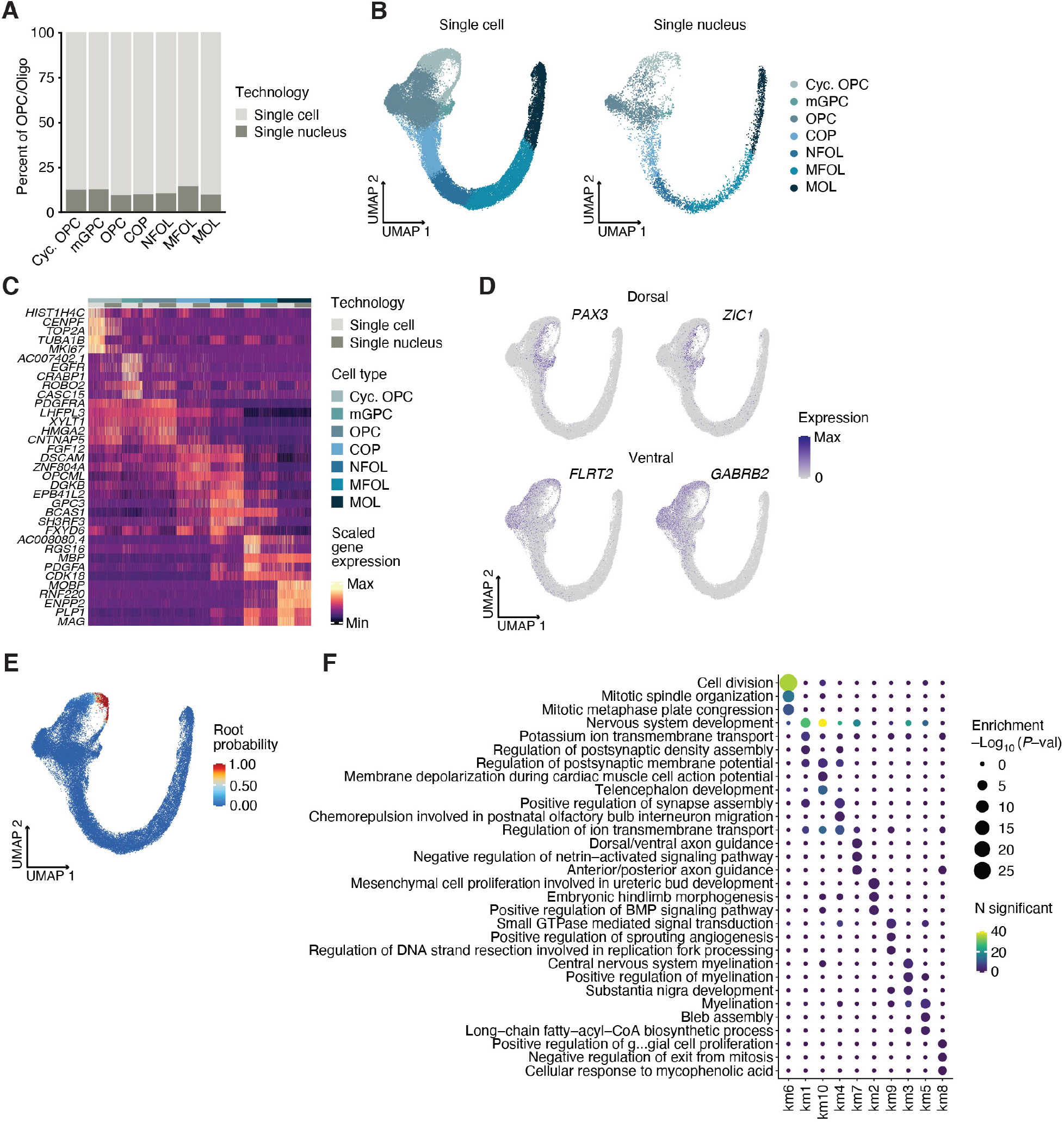
Quality control and cell types in the OPC/Oligo lineage. **(A)** Bar plot showing the percent of single cells and single nuclei in the OPC/Oligo subcluster. **(B)** UMAP of the OPC/Oligo subcluster split to show single cell and single nucleus samples separately. **(C)** Heatmap showing scaled gene expression of the top 5 genes per cell type in the OPC/Oligo subcluster (see also **Table S6**). **(D)** UMAP plots showing expression of genes with dorsal or ventral identity in the OPC/Oligo subcluster. **(E)** Root probability in UMAP space as computed by scVelo. **(F)** Bubble plot showing Gene Ontology (GO) enrichment analysis of genes represented in the 10 clusters of velocity variable genes in **Figure 2D**. Enrichment was computed using the ‘TopGO’ R package.

**Supplementary Figure 4:**
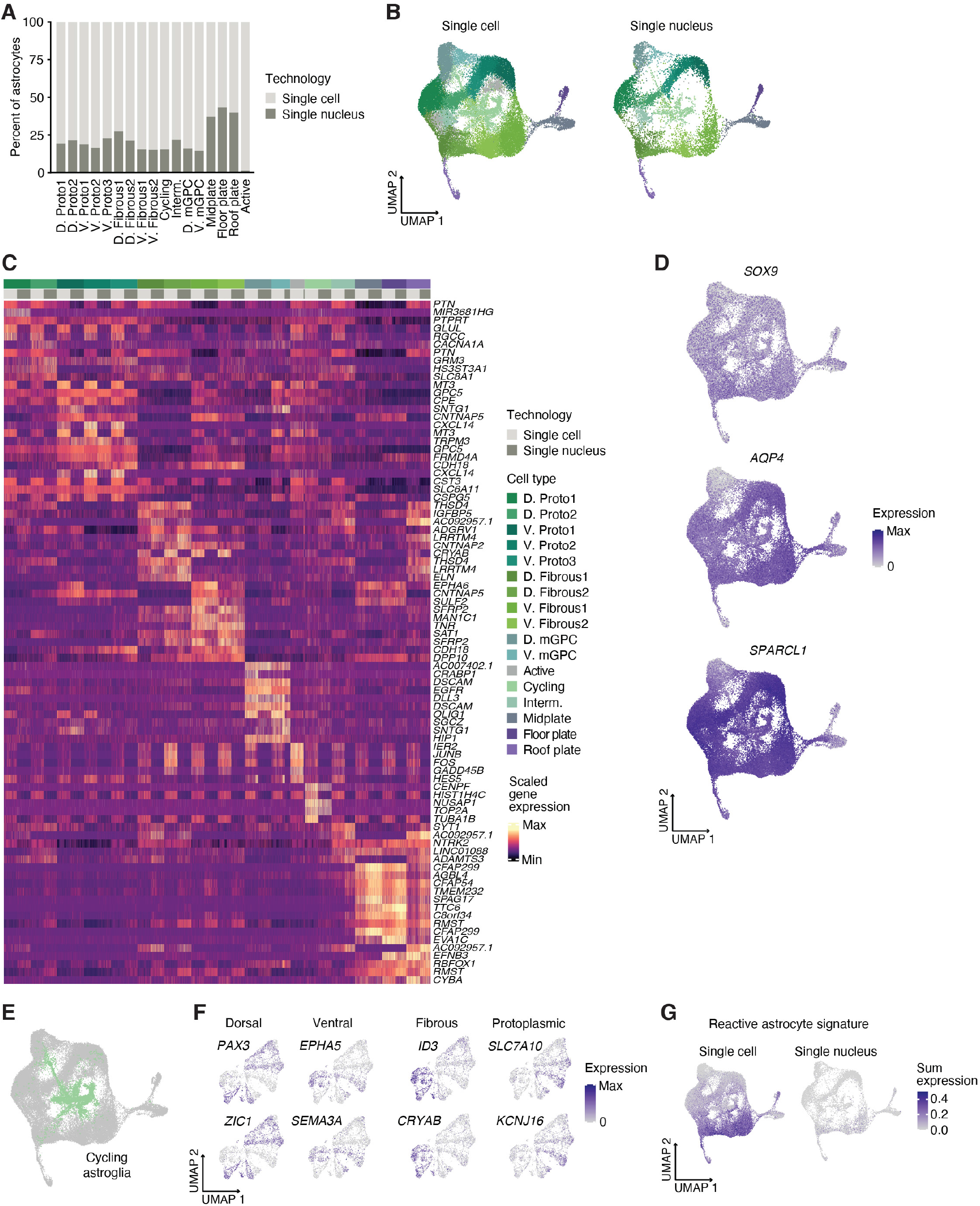
Cell types and quality control in the astroglia subcluster. **(A)** Bar plot showing the percent of single cells and single nuclei in the astroglia subcluster. **(B)** UMAP of astroglia subcluster split to show single cell and single nucleus samples separately. **(C)** Heatmap showing scaled gene expression of the top 5 non-unique genes per cell type in the astroglia subcluster (see also **Table S8**). **(D)** UMAP plots showing expression of astroglia genes. **(E)** Highlight of cycling astroglia, highlighting the cells selected for subclustering. **(F)** UMAP plots showing gene expression of markers associated with dorsal and ventral, and fibrous and protoplasmic identities within the cycling astroglia subcluster. **(G)** Reactive astrocyte signatures shown as sum expression of reactive astrocyte genes from (Liddelow et al., 2017) in single cell and single nucleus samples separately.

**Supplementary Figure 5:**
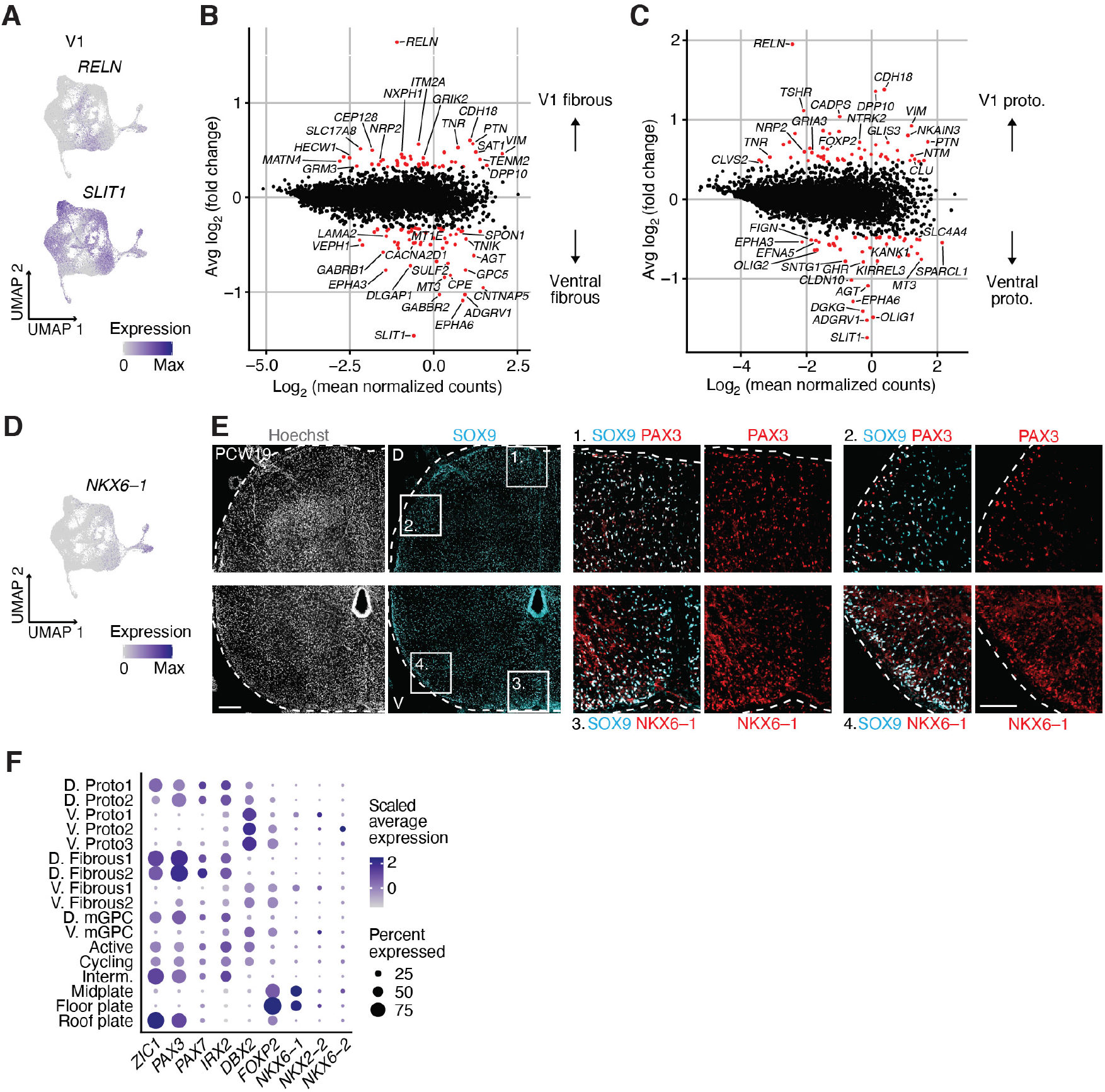
Positional identity signatures in spinal cord astrocytes. **(A)** UMAP plots showing gene expression of V1 astrocyte-associated markers. **(B)** MA plot showing differential expression between V1 fibrous astrocytes and the rest of ventral fibrous astrocytes. V1 fibrous astrocytes were selected based on their expression of *RELN* and absence of *SLIT1* (*RELN* counts > 1, *SLIT1* counts = 0). Red dots indicate genes in the top one percent of differentially expressed genes by log fold change. **(C)** MA plot showing differential expression between V1 protoplasmic astrocytes and the rest of ventral protoplasmic astrocytes. V1 protoplasmic astrocytes were selected based on their expression of *RELN* and absence of *SLIT1* (*RELN* counts > 1, *SLIT1* counts = 0). Red dots indicate genes in the top one percent of differentially expressed genes by log fold change. **(D)** UMAP plot showing expression of *NKX6-1* in the astroglia subcluster. **(E)** Representative immunohistochemistry images of SOX9 astrocytes expressing either the dorsal marker PAX3 or the ventral marker NKX6-1 in a coronal PCW19 spinal cord cryosection. Top and bottom images correspond to different cryosections. **(F)** Dot plot showing the expression of transcription factors associated with patterning in the spinal cord. The size of the dots represents the percent of cells expressing each gene while the color depicts the scaled average expression per cell type. Scale bars: 100 μm (insets in **E**), 200 μm (**E**).

**Supplementary Figure 6:**
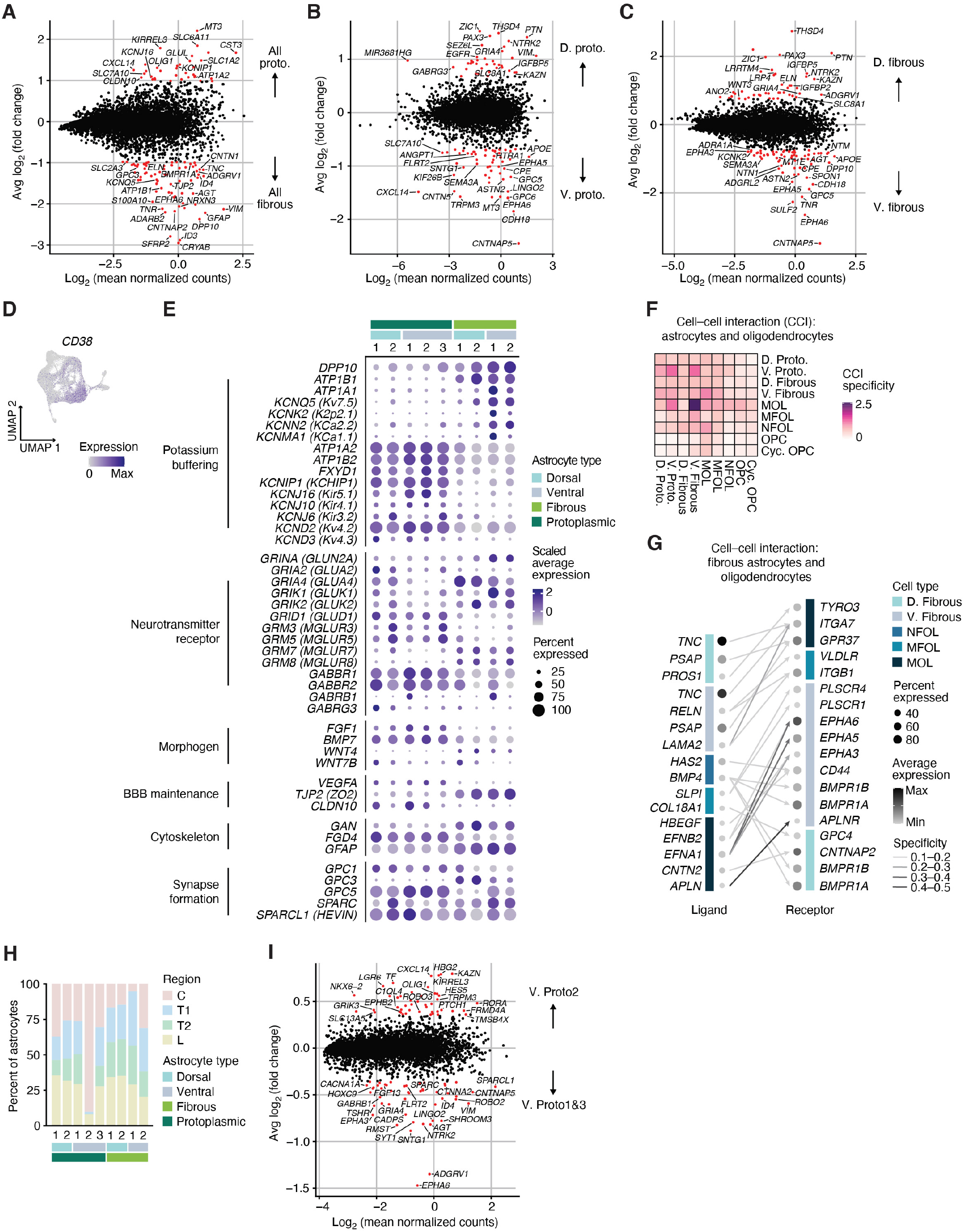
Fibrous and protoplasmic astrocytes in the spinal cord. **(A)** MA plot showing differential expression between protoplasmic and fibrous astrocytes. Red dots indicate genes in the top one percent of differentially expressed genes by log fold change. **(B)** MA plot showing differential expression between dorsal and ventral protoplasmic astrocytes. Red dots indicate genes in the top one percent of differentially expressed genes by log fold change. **(C)** MA plot showing differential expression between dorsal and ventral fibrous astrocytes. Red dots indicate genes in the top one percent of differentially expressed genes by log fold change. **(D)** UMAP plot showing expression of *CD38* in the astroglia subcluster. **(E)** Dot plot showing the expression of selected genes associated with astrocyte functions in fibrous and protoplasmic subtypes. The size of the dots represents the percent of cells expressing each gene while the color depicts the scaled average expression per subtype. **(F)** Heatmap showing the sum of specificity scores for all interactions between fibrous and protoplasmic astrocytes and OPC/Oligo cell types as computed by NATMI (see **Methods**). **(G)** Network plot showing interactions between sender and receiver cell types for ligand and receptor pairs. Only fibrous astrocytes and late-stage OPC/Oligo cell types are shown. The size of the dots represents the percent of cells expressing each gene, while the color of the dots depicts their average expression. The color of the line represents the specificity of each interaction as computed by NATMI. **(H)** Bar plot showing fibrous and protoplasmic astrocyte clusters per rostro-caudal region in PCW18 spinal cord. **(I)** MA plot showing differential expression between the V. Proto2 astrocyte cluster and the rest of the ventral protoplasmic clusters (V. Proto 1 and 3). Red dots indicate genes in the top one percent of differentially expressed genes by log fold change.

**Supplementary Figure 7:**
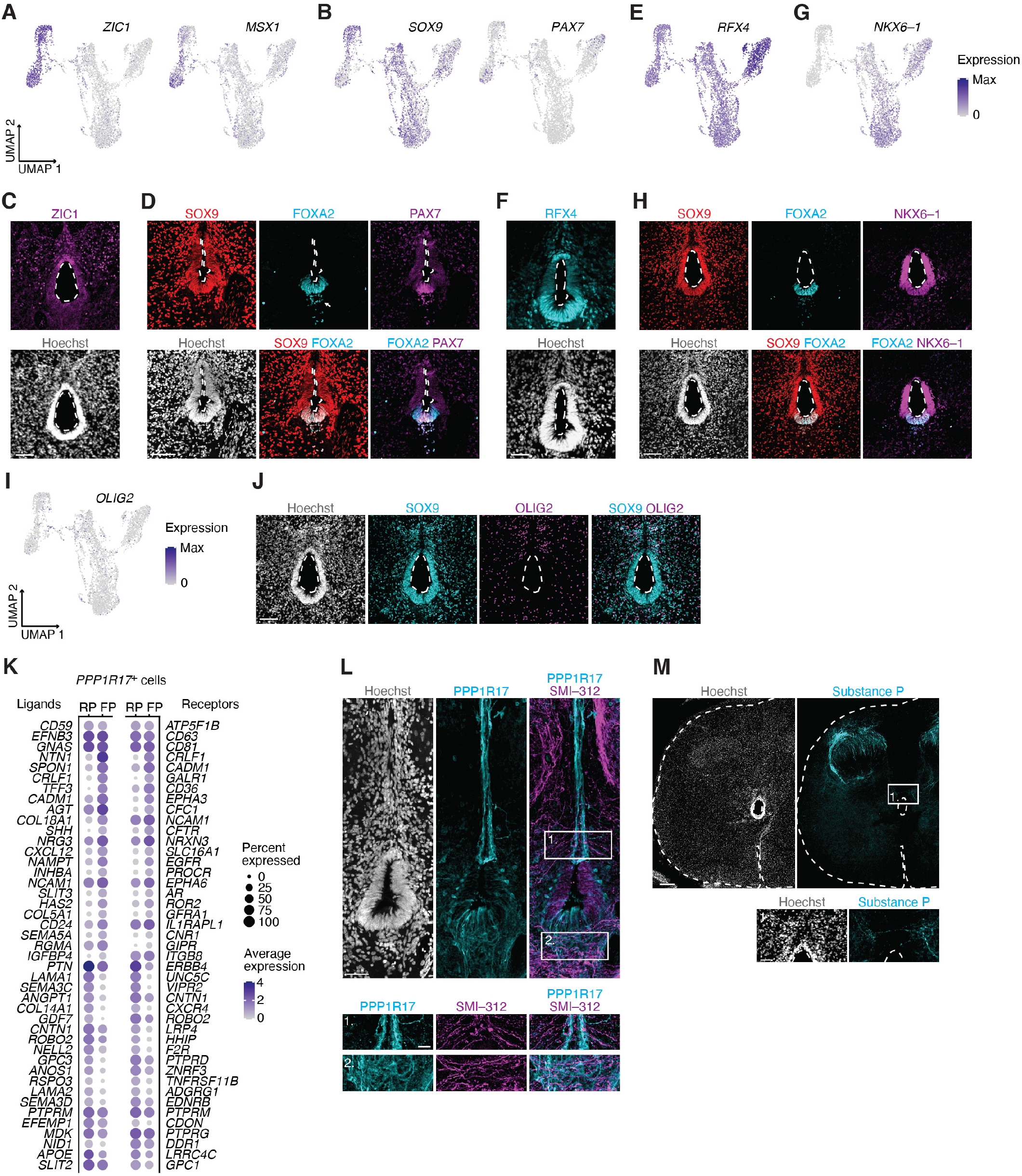
Ventricular zone (VZ) cells in the human spinal cord at midgestation. **(A)** UMAP plot showing expression of *ZIC1* in the VZ subcluster. **(B)** UMAP plots showing expression of *SOX9* and *PAX7* in the VZ subcluster. **(C)** Representative immunohistochemistry image showing ZIC1 in the VZ of PCW19 spinal cord. **(D)** Representative immunohistochemistry image showing SOX9, FOXA2 and PAX7 in the VZ of PCW19 spinal cord. Arrow shows dissociation of ventral cells from the ventricular wall. **(E)** UMAP plot showing expression of *RFX4* in the VZ subcluster. **(F)** Representative immunohistochemistry image showing RFX4 in the VZ of PCW19 spinal cord. **(G)** UMAP plot showing expression of *NKX6-1* in the VZ subcluster. **(H)** Representative immunohistochemistry image showing SOX9, FOXA2 and NKX6-1 in the VZ of PCW19 spinal cord. **(I)** UMAP plot showing expression of *OLIG2* in the VZ subcluster. **(J)** Representative immunohistochemistry image showing SOX9 and OLIG2 in the VZ of PCW19 spinal cord. **(K)** Bubble plot showing expression of ligands and receptors in *PPP1R17*^+^ cells of the roof plate (RP) and floor plate (FP). The first 3 genes were differentially expressed in both FP and RP cells relative to midplate cells. The next 20 were differentially expressed in FP relative to RP cells, and the final 20 were differentially expressed in RP relative to FP cells. Genes were identified as ligands or receptors based on the NATMI database. **(L)** Representative immunohistochemistry image showing sites of axon crossing (SMI-312) along the dorsal and ventral midline expressing PPP1R17. **(M)** Representative immunohistochemistry image showing substance P^+^ axons at the midline in PCW19 spinal cord. Scale bars: 20 μm (inset in **L**), 50 μm (**F**, **L**, inset in **M**), 100 μm (**C**, **D**, **H**, **J**), 200 μm (**M**).

**Supplementary Figure 8:**
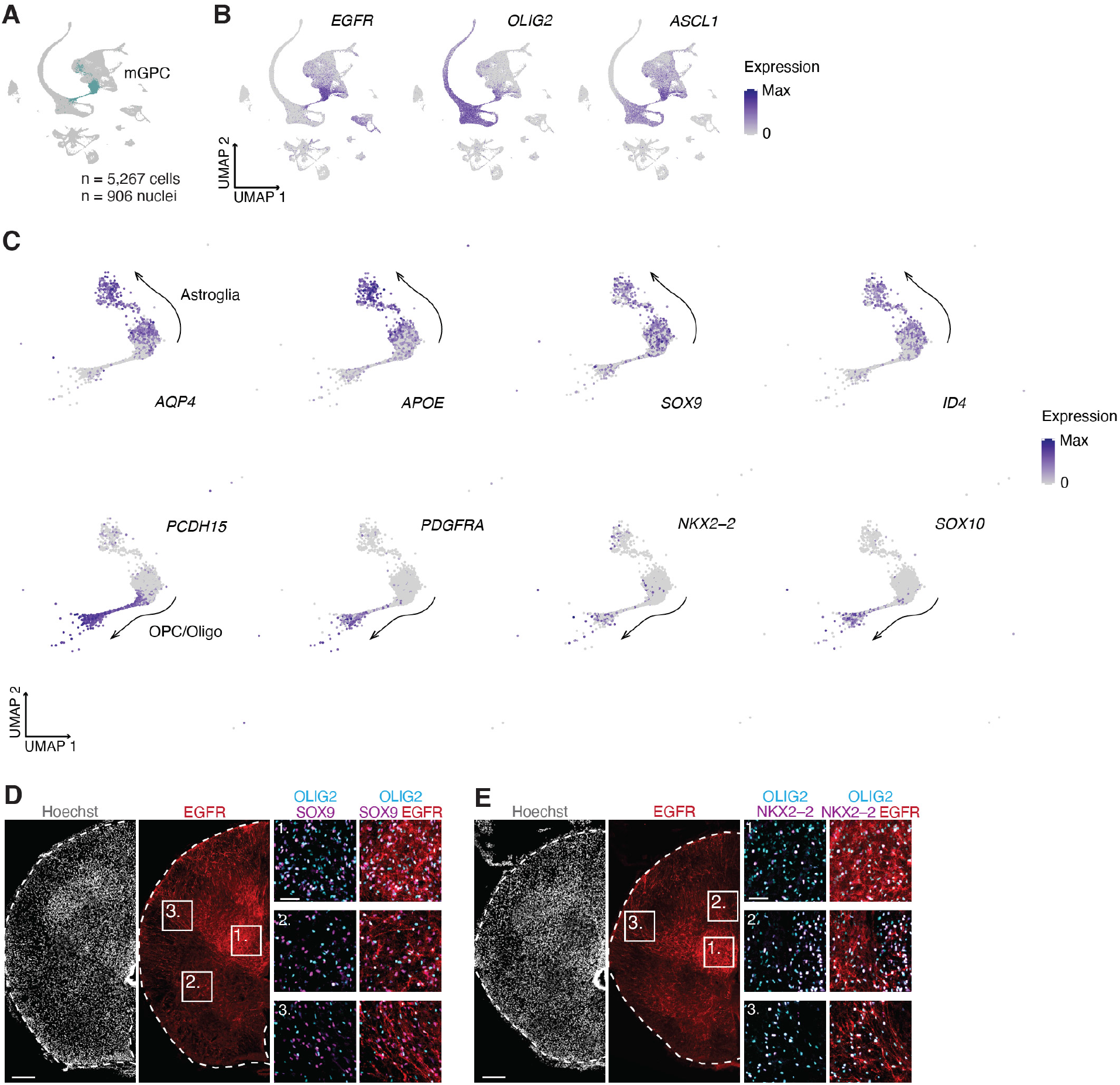
mGPCs are split into the astrocyte and oligodendrocyte lineages in the spinal cord. **(A)** Highlight of multipotent glial progenitor cells (mGPC) in the main UMAP. **(B)** UMAP plots showing expression of *EGFR, OLIG2* and *ASCL1* in the main UMAP. **(C)** UMAP plots showing expression of astroglia and OPC/Oligo genes in mGPCs within the main UMAP. **(D)** Representative immunohistochemistry images showing EGFR^+^/OLIG2^+^ mGPCs that colocalize with the astroglia-specific marker SOX9. **(E)** Representative immunohistochemistry images showing EGFR^+^/OLIG2^+^ mGPCs that colocalize with the oligodendrocyte lineage marker NKX2-2. Scale bars: 50 μm (insets in **D**, **E**), 200 μm (**D**, **E**).

**Supplementary Figure 9:**
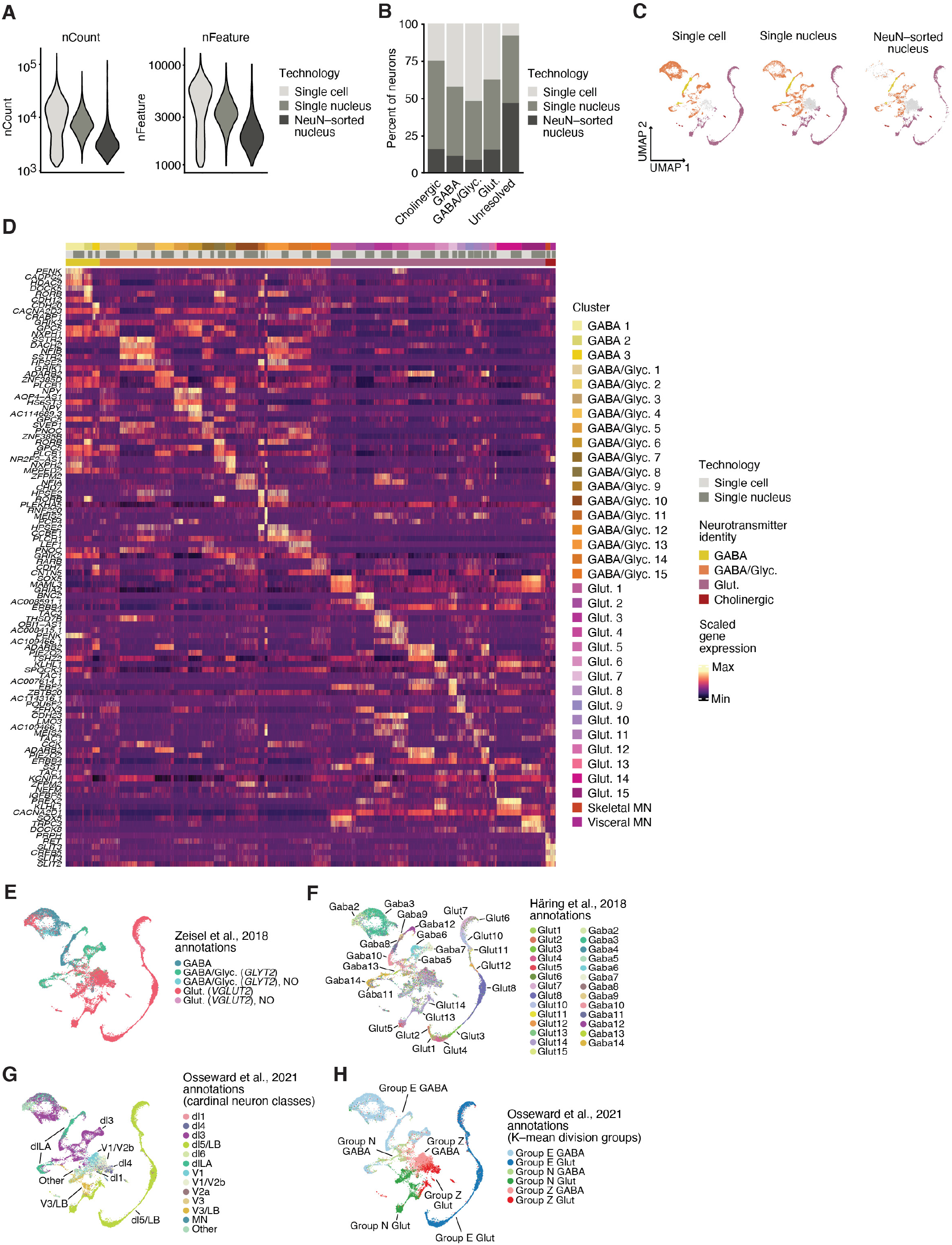
Diversity of neurons in the spinal cord. **(A)** Violin plots showing nCount (left) and nFeature (right) in the neuron subcluster separated by single cell, single nucleus and NeuN-sorted nucleus samples. **(B)** Bar plot showing the percent of single cells and single nuclei in the neuron subcluster. **(C)** UMAP of the neuron subcluster to show single cell, single nucleus and NeuN-sorted nuclei samples separately. **(D)** Heatmap showing scaled gene expression of the top 3 non-unique genes per Seurat cluster in the neuron subcluster (see also **Table S16**). **(E)** Label transfer showing neuronal annotations from Zeisel et al., 2018 in the neuron subcluster. **(F)** Label transfer showing annotations from Häring et al., 2018 in the neuron subcluster. **(G)** Label transfer showing annotations from Osseward et al., 2021 based on cardinal spinal cord neuronal classes in the neuron subcluster. **(H)** Label transfer showing annotations from Osseward et al., 2021 based on K-means division groups in the neuron subcluster. See also **Table S17** for a label transfer summary in the neuron subcluster.

**Supplementary Figure 10:**
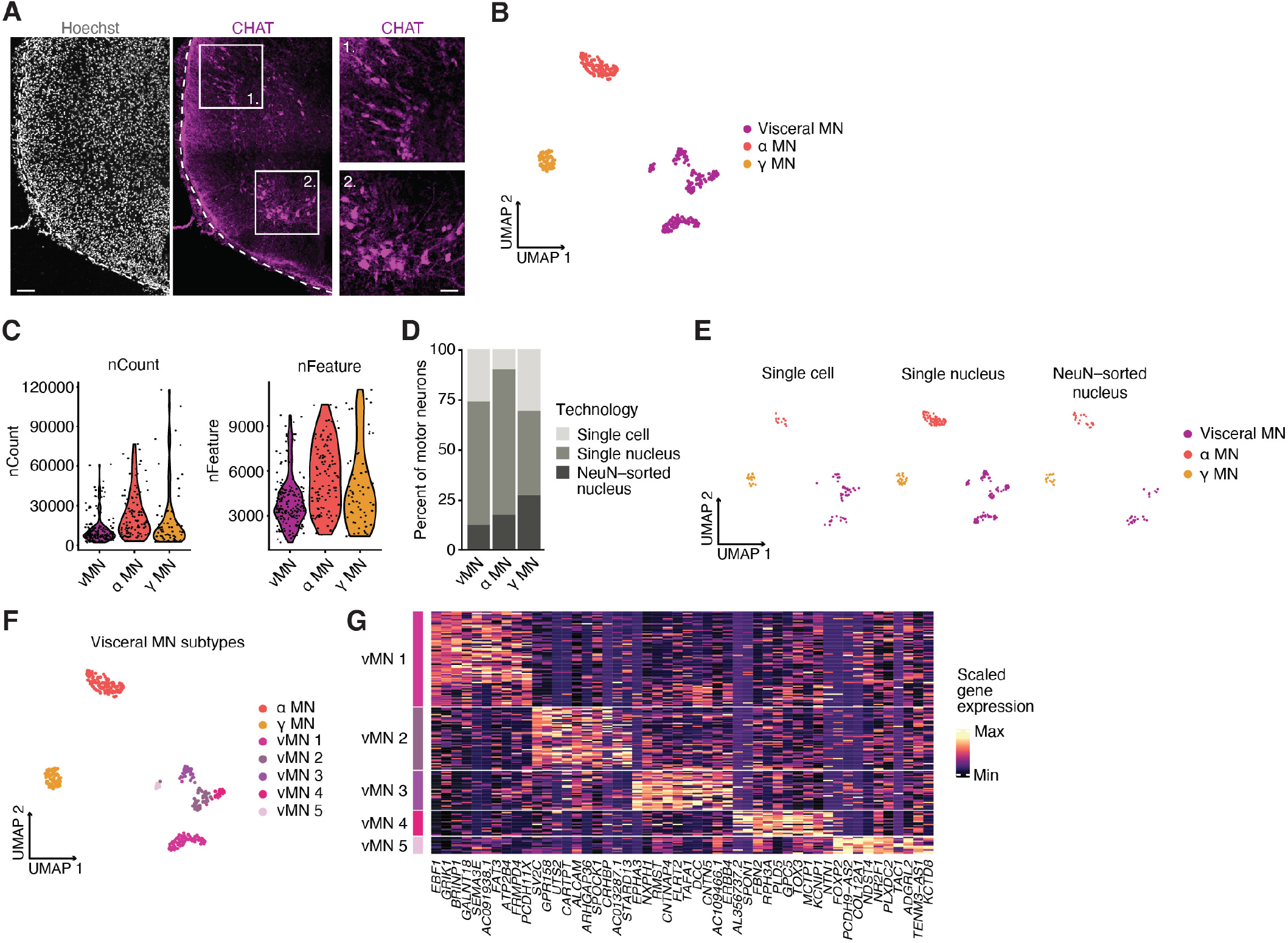
Cell types and quality control metrics in the motor neuron subcluster. **(A)** Representative immunohistochemistry images of CHAT motor neurons in a PCW19 spinal cord cryosection. **(B)** UMAP of the neuron subcluster, colored by motor neuron subtype. **(C)** Violin plots of nCount (left) and nFeature (right) separated by motor neuron subtype. **(D)** Bar plot showing the percent of single cells, single nuclei and NeuN-sorted nuclei per motor neuron subtype. **(E)** UMAP of the motor neuron subcluster split to show single cell, single nucleus and NeuN-sorted nucleus samples separately. **(F)** UMAP of the neuron subcluster showing visceral motor neuron clusters. **(G)** Heatmap showing scaled expression of the top 10 genes in visceral motor neuron clusters. Scale bars: 50 μm (insets in **A**), 100 μm (**A**).

**Supplementary Figure 11:**
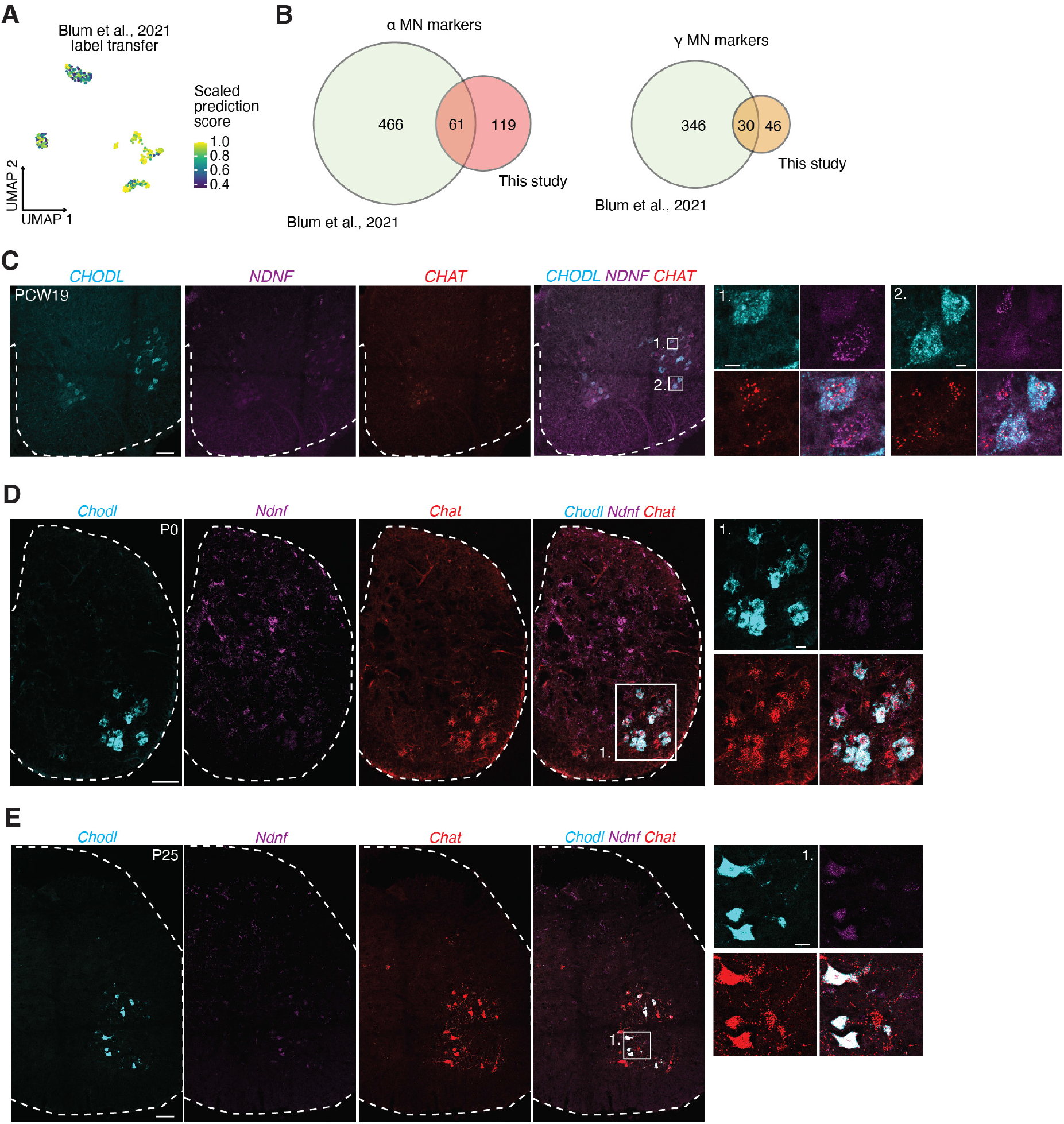
NDNF is a human-specific gamma motor neuron marker. **(A)** UMAP plot showing the scaled prediction scores for the label transfer of Blum et al., 2021 motor neuron annotations onto the motor neuron subcluster (this study). **(B)** Venn diagrams showing the number of common markers between Blum et al., 2021 and this study for alpha motor neurons (left) and gamma motor neurons (right). **(C)** Representative *in situ* hybridization of *CHAT, CHODL* and *NDNF* in coronal cervical spinal cord cryosections at PCW19. Insets show *CHAT*^+^ motor neurons that express either the alpha marker *CHODL* or the gamma marker *NDNF*. **(D)** Representative *in situ* hybridization of *Chat, Chodl* and *Ndnf* in coronal mouse spinal cord cryosections at P0. Inset shows *Chat^+^* motor neurons that express the alpha marker *Chodl* but not the gamma marker *Ndnf*. **(E)** Representative *in situ* hybridization of *Chat, Chodl* and *Ndnf* in coronal mouse spinal cord cryosections at P25. Insets show *Chat^+^* motor neurons that express both the alpha marker *Chodl* and the gamma marker *Ndnf*. Scale bars: 10 μm (insets in **C**), 20 μm (insets in **D**, **E**), 100 μm (**C**, **D**, **E**).

**Supplementary Figure 12:**
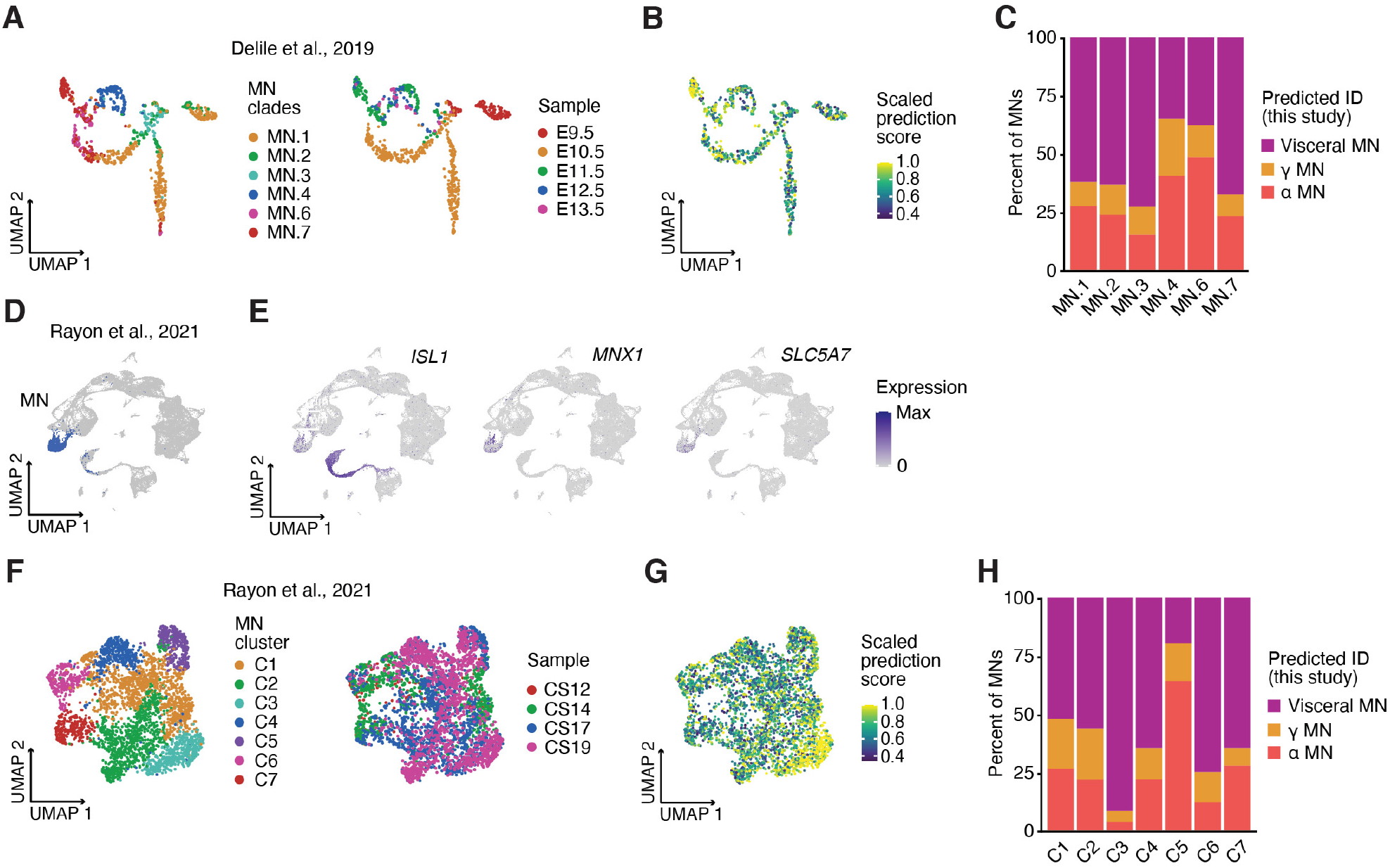
Separation of alpha and gamma motor neurons at midgestation in humans. **(A)** UMAP plots showing motor neurons in Delile et al., 2019, colored by their assigned clades (left) and sample age (right). **(B)** UMAP plot showing the scaled prediction scores for the label transfer of this study’s motor neuron type annotations onto Delile et al., 2019 motor neurons. **(C)** Bar plot showing the percent of predicted identities based on annotations from this study in Delile et al., 2019 MN clades. **(D)** UMAP plot of all cells in Rayon et al., 2021 highlighting the MN cluster selected for further analysis. **(E)** UMAP plots showing gene expression of motor neuron markers in Rayon et al., 2021. **(F)** UMAP plots showing the motor neuron subcluster in Rayon et al., 2021, colored by Seurat clusters (left) and sample age (right). Approximately, Carnegie Stage (CS) 12 corresponds to PCW4; CS14 to PCW5; CS17 to PCW6; and CS19 to PCW7. **(G)** UMAP plot showing the scaled prediction scores for the label transfer of this study’s motor neuron type annotations onto Rayon et al., 2021 motor neurons. **(H)** Bar plot showing the percent of predicted identities based on annotations from this study in Rayon et al., 2021 motor neuron clusters.

**Supplementary Figure 13:**
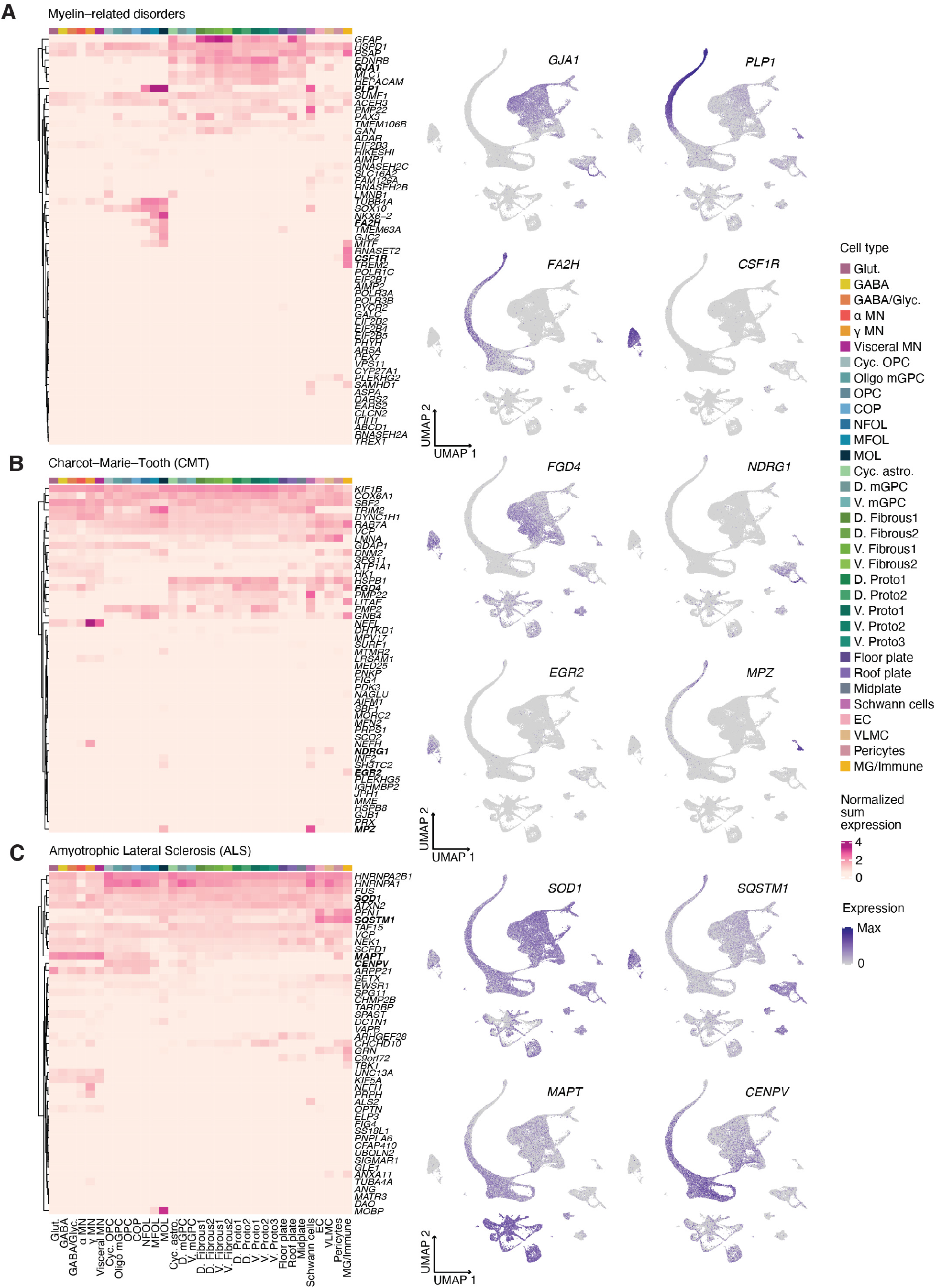
Mapping of disease gene sets on human developing spinal cord atlas. **(A)** Heatmap (left) showing the sum of gene counts normalized to total counts per cell cluster for genes associated with myelin-related disorders; and UMAP plots (right) showing gene expression of selected genes. See **Methods** for references. **(B)** Heatmap (left) showing the sum of gene counts normalized to total counts per cell cluster for genes associated with Charcot-Marie-Tooth (CMT); and UMAP plots (right) showing expression of selected genes. See **Methods** for references. **(C)** Heatmap (left) showing the sum of gene counts normalized to total counts per cell cluster for genes associated with amyotrophic lateral sclerosis (ALS); and UMAP plots (right) showing expression of selected genes. See **Methods** for references.

## List of Supplementary Tables

Supplementary Tables 1–24 were uploaded separately in Excel format, with each table in a separate tab.

| Title | Legend |
| --- | --- |
| **Supplementary Table 1.** List of all samples used in this study | Information regarding age, sample preparation, number of cells, mean reads and median number of genes is provided for each sample |
| **Supplementary Table 2.** Cell type marker genes in single cell samples | Marker genes were filtered by adjusted p-value < 0.01 and avg log<sub>2</sub>FC > 0.1 |
| **Supplementary Table 3.** Cell type marker genes in single nuclei samples | Marker genes were filtered by p-value < 0.01 and avg log<sub>2</sub>FC > 0.1 |
| **Supplementary Table 4.** GO terms of genes enriched in single cell and single nucleus samples | Top 5 Gene Ontology (GO) terms of genes enriched in either single cell or single nucleus samples with the genes represented in each term. Cells and nuclei of the same type were compared to one another via differential expression. The top 250 genes were then selected by p-value < 0.01 and logFC > 0 or logFC < 0 and enriched using a hypergeometric test. |
| **Supplementary Table 5.** Cell type marker genes in integrated single cell and single nucleus samples | Marker genes were filtered by adjusted p-value < 0.01 and avg log<sub>2</sub>FC > 0.1 |
| **Supplementary Table 6.** Cell type marker genes in OPC/Oligo subcluster | Marker genes were filtered by adjusted p-value < 0.01 and avg log<sub>2</sub>FC > 0.1 |
| **Supplementary Table 7.** Genes with high velocities in the OPC/Oligo subcluster | The top 20 genes for each cell type have been identified using the scVelo function 'rank_velocity_genes' |
| **Supplementary Table 8.** Cell type marker genes in the astroglia subcluster | Marker genes were filtered by adjusted p-value < 0.01 and avg log<sub>2</sub>FC > 0.1 |
| **Supplementary Table 9.** Differentially expressed genes in protoplasmic vs fibrous astrocytes | Marker genes were filtered by adjusted p-value < 0.01 and avg log<sub>2</sub>FC > 0.1 |
| **Supplementary Table 10.** Differentially expressed genes in dorsal vs ventral fibrous astrocytes | Marker genes were filtered by adjusted p-value < 0.01 and avg log<sub>2</sub>FC > 0.1 |
| **Supplementary Table 11.** Differentially expressed genes in dorsal vs ventral protoplasmic astrocytes | Marker genes were filtered by adjusted p-value < 0.01 and avg log<sub>2</sub>FC > 0.1 |
| **Supplementary Table 12.** Cell-cell interaction network edges | Network edges as calculated by NATMI for interactions between fibrous and protoplasmic astrocytes and OPC/Oligo cell types |
| **Supplementary Table 13.** Cell type marker genes in VZ subcluster | Marker genes were filtered by adjusted p-value < 0.01 and avg log<sub>2</sub>FC > 0.1 |
| **Supplementary Table 14.** Differentially expressed genes in floor plate *PPP1R17* positive vs negative cells | Cells were filtered based on *PPP1R17* counts > 0. Marker genes were filtered by adjusted p-value < 0.01 and avg log<sub>2</sub>FC > 0.1 |
| **Supplementary Table 15.** Differentially expressed genes in roof plate *PPP1R17* positive vs negative cells | Cells were filtered based on *PPP1R17* counts > 0. Marker genes were filtered by adjusted p-value < 0.01 and avg log<sub>2</sub>FC > 0.1 |
| **Supplementary Table 16.** Cell cluster marker genes in neuron subcluster | Marker genes were filtered by adjusted p-value < 0.01 and avg log<sub>2</sub>FC > 0.1 |
| **Supplementary Table 17.** Label transfer summary for neuron subcluster | Summary of label transfer results between neuron clusters in this study and reference datasets. For each Seurat cluster in this study, the proportion of cells matching the most common label of the cluster and the average Seurat robustness score are shown. |
| **Supplementary Table 18.** Differential expressed genes in skeletal vs visceral MNs | Marker genes were filtered by adjusted p-value < 0.01 and avg log<sub>2</sub>FC > 0.1 |
| **Supplementary Table 19.** Cell cluster marker genes in visceral MNs | Marker genes were filtered by adjusted p-value < 0.01 and avg log<sub>2</sub>FC > 0.1 |
| **Supplementary Table 20.** Differentially expressed genes in alpha vs gamma MNs | Marker genes were filtered by adjusted p-value < 0.01 and avg log<sub>2</sub>FC > 0.1 |
| **Supplementary Table 21.** Differentially expressed genes in alpha MNs in this study vs Blum et al., 2021 | Marker genes were filtered by adjusted p-value < 0.01 and avg log<sub>2</sub>FC > 0.1 |
| **Supplementary Table 22.** Differentially expressed genes in gamma MNs in this study vs Blum et al., 2021 | Marker genes were filtered by adjusted p-value < 0.01 and avg log<sub>2</sub>FC > 0.1 |
| **Supplementary Table 23.** List of disease-associated genes | List of disease-associated genes used for **Extended Data Fig. 13** with their corresponding references |
| **Supplementary Table 24.** List of antibodies used in this study | For each primary antibody used for immunohistochemistry, the host species, dilution, company and catalogue number are provided |

